# A Consensus Framework Unifies Multi-Drug Synergy Metrics

**DOI:** 10.1101/683433

**Authors:** David J. Wooten, Christian T. Meyer, Vito Quaranta, Carlos Lopez

**Author notes:** These authors contributed equally to this work (David J. Wooten, Christian T. Meyer). Correspondence should be directed to C.F.L.

## Abstract

Drug combination discovery depends on reliable synergy metrics; however, no consensus exists on the appropriate synergy model to prioritize lead candidates. The fragmented state of the field confounds analysis, reproducibility, and clinical translation of combinations. Here we present a mass-action based formalism to accurately measure the synergy of drug combinations. In this work, we clarify the relationship between the dominant drug synergy principles and show how biases emerge due to intrinsic assumptions which hinder their broad applicability. We further present a mapping of commonly used frameworks onto a unified synergy landscape, which identifies fundamental issues impacting the interpretation of synergy in discovery efforts. Specifically, we infer how traditional metrics mask consequential synergistic interactions, and contain biases dependent on the Hill-slope and maximal effect of single-drugs. We show how these biases systematically impact the classification of synergy in large combination screens misleading discovery efforts. The proposed approach has potential to accelerate the translatability and reproducibility of drug-synergy studies, by bridging the gap between the curative potential of drug mixtures and the complexity in their study.

## Introduction

Throughout the preceding century, two dominant principles have been used to quantify synergy of drug combinations: the Dose Equivalence Principle (DEP), introduced by Loewe [1, 2] and expanded by Chou [3], and the Multiplicative Survival Principle (MSP), introduced by Bliss [4]. Discrepancies between these two principles, however, have resulted in divergent conclusions between synergy studies [5]. In 1992, a committee was convened in Saariselkä, Finland seeking a consensus between these principles [6, 7]. However, unable to reconcile their differences, the committee’s conclusion, codified as The Saariselkä Agreement, simply recommended drug combination studies explicitly state how synergy was calculated [6, 7].

More recently, a proliferation of synergy models, derived as extensions of either the DEP or MSP, has further splintered the field [8, 9, 10, 11, 12, 13]. In the absence of a consensus framework for drug synergy, discovery efforts for combinations often calculate all available synergy metrics [14, 15, 16], as first recommended by Greco and colleagues [5] following Saariselkä. However, there remains no basis for choosing one metric over another, which becomes particularly problematic when synergy metrics are in conflict. This “calculate everything” paradigm thus hampers reproducibility between studies, delays progress in the discovery of truly synergistic drug combinations, and negatively impacts the translatability of combination discovery efforts.

Despite the lack of consensus on how to quantify synergy, drug combination screens remain essential to both pharmaceutical and academic discovery efforts, as shown in recent challenges by AstraZeneca and the NCI-DREAM consortia [17, 18], as well as combinatorial CRISPR screens [19]. Yet, the paucity of successful clinical combinations explicable by true pharmacological interaction, rather than patient-to-patient variability [20], is symptomatic of the challenges facing the field. Therefore, the need identified at Saariselkä still exists: a consensus framework to interpret drug combination pharmacology.

We recently introduced a new model to quantify synergy based on the Law of Mass Action, named Multi-dimensional Synergy of Combinations (MuSyC) [21]. In this work, we show that our theory generalizes the DEP and MSP, thereby unifying the field of drug synergy, as sought at Saariselkä. We further map the landscape of current synergy metrics, including: Bliss Independence [4], Loewe Additivity [1], Combination Index (CI) [3], Highest Single Agent (HSA) [22], Effective Dose model by Zimmer et al. [8], ZIP [9], a partial differential equation (PDE) Hill model by Schindler [10], and BRAID [13]. In mapping relationships between these various metrics, we identified systematic differences impacting the interpretation of synergy in drug combination experiments. Specifically, we found: (1) the conflation of synergistic potency and efficacy masks synergistic interactions; (2) MSP frameworks are biased toward antagonism for drugs with intermediate efficacy; and (3) DEP frameworks contain a Hill-slope dependent bias. The Hill-slope bias results from satisfying the famous “sham” combination thought experiment, arguing against the merit of sham-compliance as a measure of validity for synergy frameworks. Using two large combination datasets [23, 24], MuSyC identifies real-world examples where the conflicting assumptions of previous drug synergy frameworks misleads or impedes drug discovery efforts through these pervasive and predictable biases. We therefore propose MuSyC as a consensus framework to interpret combination pharmacology and signify its broad applicability to the study of drug mixtures.

### A multi-dimensional formalism to measure multi-drug synergistic effects

The 4-parameter Hill equation is commonly used to fit dose-response data from in *vitro* and *in vivo* assays (see Box 1 equation (3) for derivation and Table 1 for parameter annotation). This equation can be derived from the equilibrium of a 2-state model of drug effect based on the Law of Mass Action (Figure 1A left). Traditionally, the parameters of the Hill equation are interpreted as a drug’s efficacy (*E*_0_ − *E*_1_), potency (*C*), and cooperativity (*h*), also known as the Hill slope. These parameters correspond to three possible geometric transformations of a dose-response curve (Figure 1A right). To generalize this one-drug formalism to two concurrent drugs, we recently developed a 4-state mass-action model of combination pharmacology (Figure 1B left) [21]. From this model, we derived a two-dimensional (2D) Hill equation for two drugs (Box 1 equation 8) defining a dose-response surface (Figure 1B middle). The 2D Hill equation contains five additional parameters, not present in the single-drug Hill equation, which measure different types of drug interactions. These additional parameters measure changes in a drug’s efficacy (*β*), potency (*α*_12_ and *α*_21_), and cooperativity (*γ*_12_ and *γ*_21_) in a combination —representing three distinct types of synergy (Figure 1B right, Table 1). As we show below, these parameters are conflated in traditional synergy metrics obscuring the true origin and magnitude of drug synergy or antagonism.

**Figure 1:**
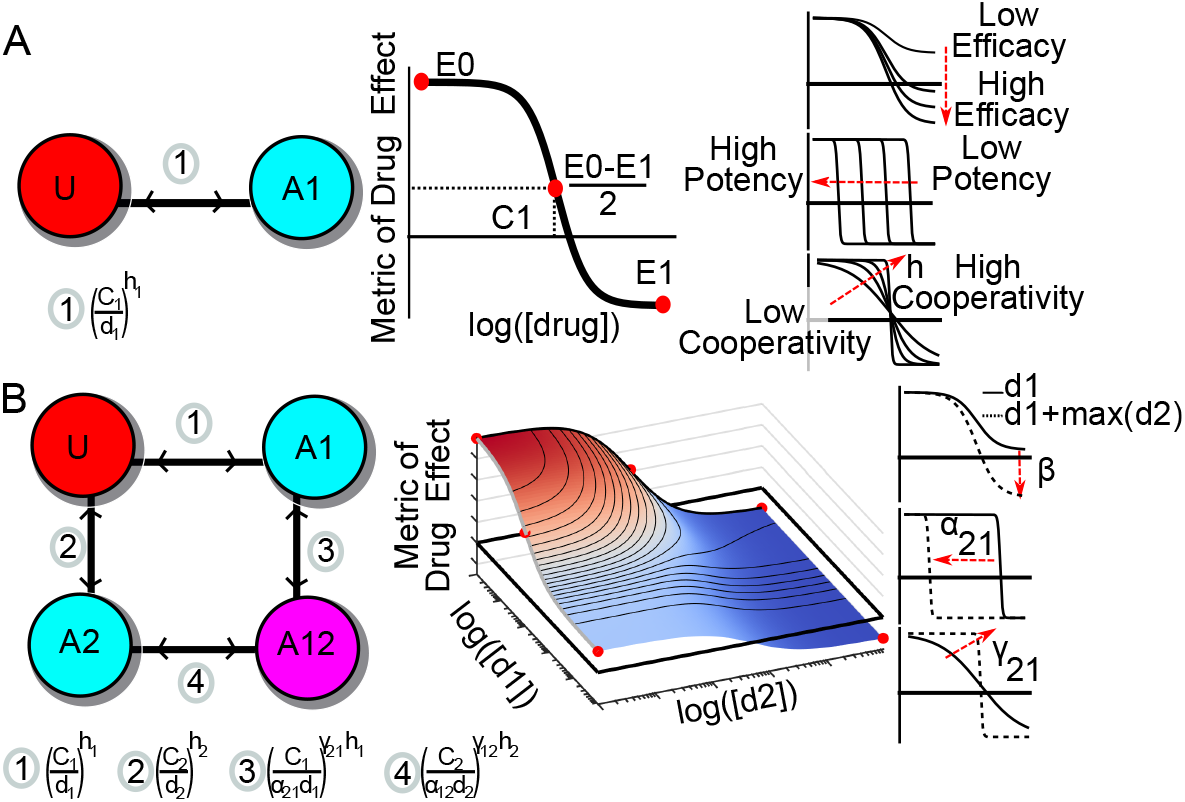
MuSyC mass action model of drug combination synergy. A) Single-drug model: The traditional equation for fitting dose-response relationships (middle) is the 4-parameter Hill equation. This equation can be derived using the Law of Mass Action from a two state model of drug effect (left). Edge notation is equal to the ratio of states at equilibrium 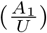. The Hill equation contains parameters measuring a drug’s efficacy (*E*_0_ − *E*_1_), potency (*C*), and cooperativity (h). Each parameter corresponds to distinct geometric transformations of the dose-response curve (right). B) Two-drug model: MuSyC is derived from a four-state mass-action model of combination pharmacology (left) and results in a 2D Hill-like equation describing a dose-response surface (middle). Edge notation denotes the ratio of the connected corners for the boundary condition. For example, edge #3 annotation means 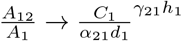 when *d*_2_ → inf. Beyond the parameters of the single Hill equation, the 2D Hill equation has additional parameters (*β, α, γ*) corresponding to distinct transformations of the dose-response surface (right) (Video S1). These transformations directly measure the changes in a single drugs’ efficacy, potency, and cooperativity due to the combination, and, therefore, are interpreted as synergistic efficacy (*β*), synergistic potency (*α*), and synergistic cooperativity (*γ*). There are two values for *α* and *γ* because each drug can independently modulate the potency and cooperativity of the other [8, 9] (edge 3 vs. edge 4 of the state transition model). In contrast, the single *β* parameter describes the percent increase in maximal effect due to both drugs (effect at A12). See Figure S1 for MuSyC extension to three drugs.

**Table 1:**
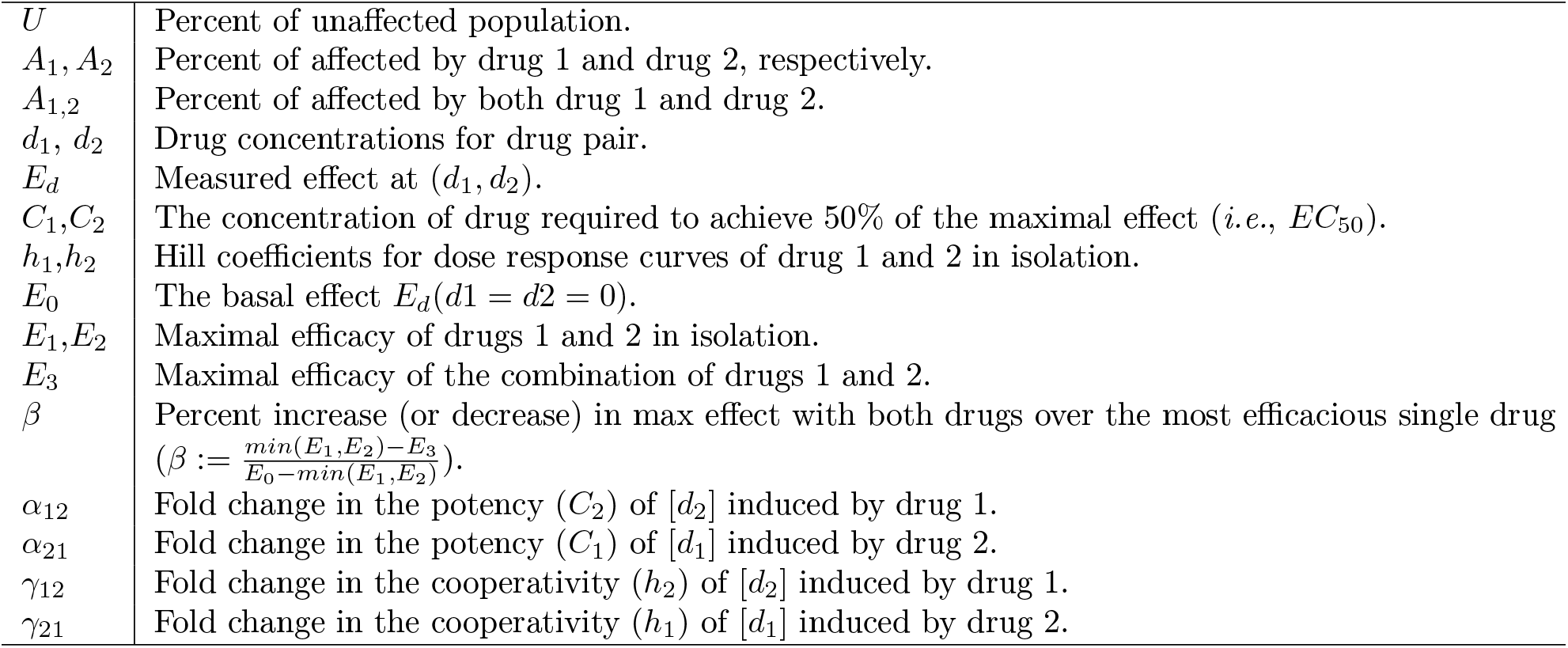
Annotation of MuSyC parameters.

|  |  |
| --- | --- |
| $U$ | Percent of unaffected population. |
| $A_1, A_2$ | Percent of affected by drug 1 and drug 2, respectively. |
| $A_{1,2}$ | Percent of affected by both drug 1 and drug 2. |
| $d_1, d_2$ | Drug concentrations for drug pair. |
| $E_d$ | Measured effect at $(d_1, d_2)$. |
| $C_1, C_2$ | The concentration of drug required to achieve 50% of the maximal effect (*i.e.*, $EC_{50}$). |
| $h_1, h_2$ | Hill coefficients for dose response curves of drug 1 and 2 in isolation. |
| $E_0$ | The basal effect $E_d(d_1 = d_2 = 0)$. |
| $E_1, E_2$ | Maximal efficacy of drugs 1 and 2 in isolation. |
| $E_3$ | Maximal efficacy of the combination of drugs 1 and 2. |
| $\beta$ | Percent increase (or decrease) in max effect with both drugs over the most efficacious single drug<br>($\beta := \frac{\min(E_1, E_2) - E_3}{E_0 - \min(E_1, E_2)}$). |
| $\alpha_{12}$ | Fold change in the potency ($C_2$) of $[d_2]$ induced by drug 1. |
| $\alpha_{21}$ | Fold change in the potency ($C_1$) of $[d_1]$ induced by drug 2. |
| $\gamma_{12}$ | Fold change in the cooperativity ($h_2$) of $[d_2]$ induced by drug 1. |
| $\gamma_{21}$ | Fold change in the cooperativity ($h_1$) of $[d_1]$ induced by drug 2. |

### A bridge between DEP and MSP maps existing synergy approaches onto a common landscape

In recent years, several alternative synergy models have been proposed. Broadly, these models are derived from one of two guiding principles: the Multiplicative Survival Principle (MSP) and the Drug Equivalence Principle (DEP) (Table 3). Prior work has shown contradictory results when comparing between MSP and DEP frameworks [5], and there remains a lack of consensus on the commonality between the two principles [6, 9, 13, 11]. Here we show MuSyC satisfies both the DEP and MSP under certain conditions (Figure 2A,B), thereby unifying the foundational principles of drug synergy.

**Figure 2:**
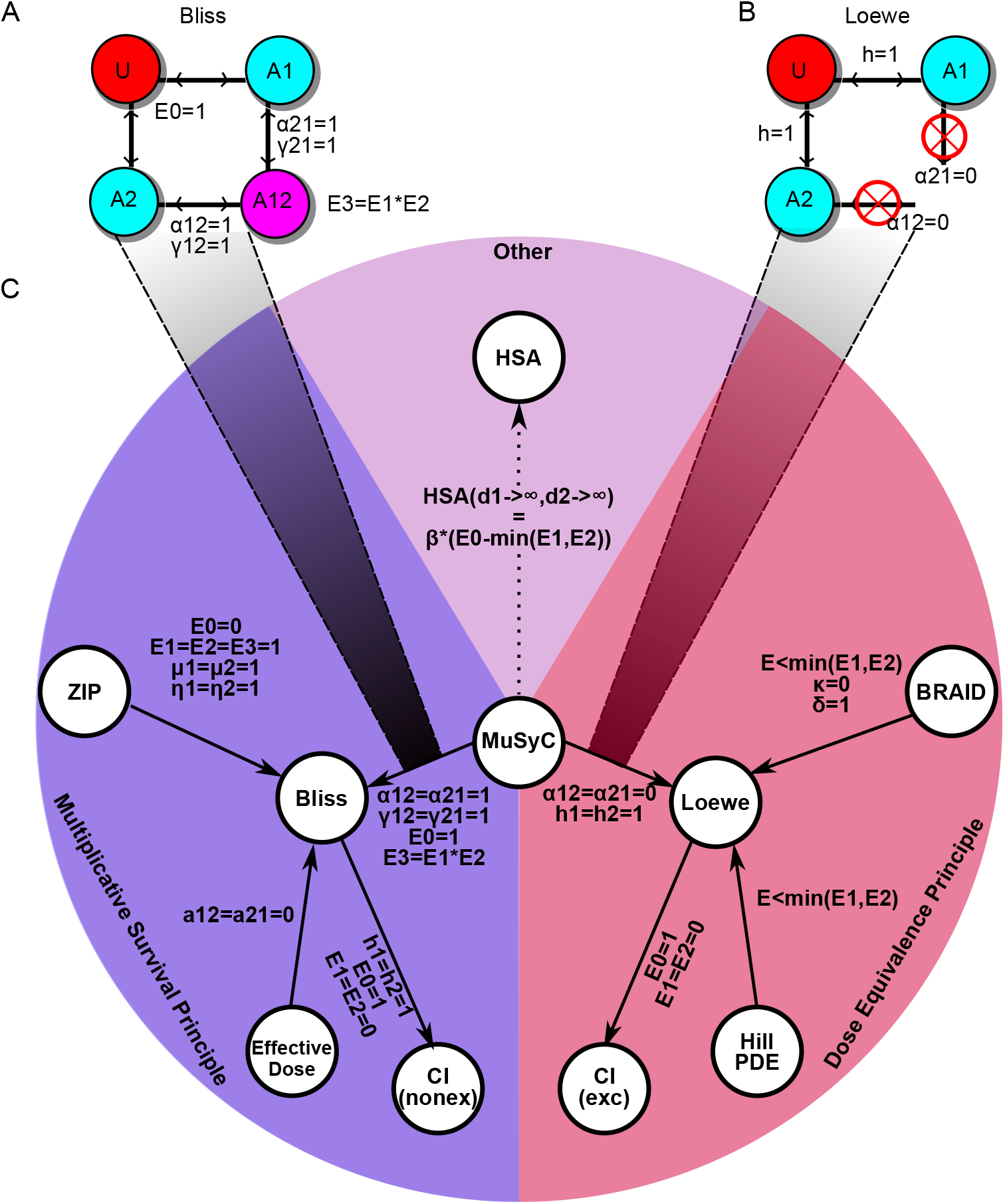
Unifying MSP and DEP with MuSyC, and mapping the landscape of drug synergy. A) The Bliss null model, the base model for all MSP frameworks, emerges from MuSyC when *E*_0_ = 1, *α*_12_ = *α*_21_ = *γ*_12_ = *γ*_21_ = 1, and *E*_3_ = *E*_1_*E*_2_. B) The Loewe null model, the base model for all DEP frameworks, emerges from MuSyC when *h*_1_ = *h*_2_ = 1 and *α*_12_ = *α*_21_ = 0. The constraint on *α* indicates the drugs’ activities are mutually exclusive (i.e., the double-drugged state *A*_1,2_ does not exist). C) Network of relationships between synergy frameworks (nodes) grouped by their underlying principle (colors). The notation next to solid edges signifies conditions under which source model reduces to end model’s null model. The dotted edge indicates MuSyC synergistic efficacy (*β*) is proportional to HSA as *d*_1_ → ∞, *d*_2_ → ∞. See Supplemental Section *Derivation of the theoretical relationships between different synergy frameworks* for complete annotation of the parameters defined in each method. Where possible, parameters from each framework were translated in terms of the dose-response parameters defined for MuSyC (Table 1) to facilitate comparison.

The MSP was first described by Bliss [4] and is the foundation of the Bliss Independence framework. MSP assumes the probability of a cell surviving treatment by drug 1 (*U*_1_) is independent of the probability of the same cell surviving treatment by drug 2 (*U*_2_). Therefore, the probability of surviving both Drug 1 and Drug 2 is *U* = *U*_1_ · *U*_2_ [4]. Synergy or antagonism occur when *U* = *U*_1_ · *U*_2_. MuSyC satisfies the MSP under the following conditions: (1) the effect metric is expressed as a percent (*E*_0_ = 1, and *E*_3_ = *E*_1_*E*_2_), (2) there is no potency synergy (*α*_12_ = *α*_21_ = 1), and (3) there is and no cooperativity synergy (*γ*_12_ = *γ*_21_ = 1) (Figure 2A, see Supplemental Section *Multiplicative Survival Principle* for details).

The DEP was first established by Loewe [1, 2] and subsequently expanded by Chou and Talalay [25]. This approach assumes that for a given effect *E*—achievable either by dose *d*_1_ of Drug 1 alone, or dose *d*_2_ of Drug 2 alone—there is a constant ratio 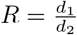 such that using Δ*d*_2_ less of Drug 2 can always be compensated for with Δ*d*_1_ = *R*Δ*d*_2_ more of Drug 1 to achieve the same effect [11]. This definition leads to the linear isoboles (contours of equal effect) characteristic of the Loewe null model (Figure S2). Synergy occurs when less one drug is required to compensate for a decrease in the other than expected by DEP. MuSyC satisfies the DEP under the following conditions: (1) the drugs are mutually exclusive (a_12_ = a_21_=0) and (2) *h_1_ = h_2_* = 1 (Figure 2B, see Supplemental Section *Dose Equivalence Principle* for details).

From the literature, we identified several prominent synergy models beyond Bliss and Loewe including: CI [3], HSA [22], Effective Dose model [8], ZIP [9], and Hill PDE [10]. Table 2 compares key features and assumption of the different synergy models. Each of these methods, as well as MuSyC, defines synergy based on the experimental deviation from a null (additive) dose-response surface. Because almost all synergy frameworks are founded on either the DEP or MSP, we standardized relationships between the various models, mapping the global landscape of drug synergy (Figure 2C, see Supplemental Section *Relationships between different synergy frameworks* for details).

**Table 2:** Comparison of traditional and modern frameworks for calculating synergy. *CI has 2 equations for synergy in the original derivation [3], for the mutually exclusive and mutually non-exclusive case. The mutually exclusive case, which is equivalent to Loewe, has been widely adopted and is the model compared here.

|  | Dose Equivalence Principle |  |  |  | Multiplicative Survival Principle |  |  | Other | Both |
| --- | --- | --- | --- | --- | --- | --- | --- | --- | --- |
|  | Loewe | BRAID | Hill PDE | Combination Index* | Bliss | Equivalent Dose | ZIP | HSA | MuSyC |
| Defined for arbitrary metrics of drug effect (not just percent data) | ✓ | ✓ |  |  |  |  | ✓ | ✓ | ✓ |
| Does not require all drugs to be equally efficacious |  | ✓ | ✓ |  | ✓ |  |  | ✓ | ✓ |
| Synergy is concentration independent |  | ✓ |  |  |  | ✓ |  |  | ✓ |
| Synergy is related to traditional dose-response parameters |  |  |  |  |  | ✓ |  | ✓ | ✓ |
| Satisfies the sham experiment | ✓ | ✓ | ✓ | ✓ |  |  |  |  |  |

In deriving this map, we uncovered potential sources of error when using MSP or DEP methods which impact interpretation of synergy studies. Specifically, we identified three recurrent considerations meriting attention from the field. 1) Previous synergy metrics conflate different synergy types (*i.e*. potency, efficacy, cooperativity) in ways that can mask synergistic and antagonistic interactions. 2) The connection between MuSyC and the MSP-derived frameworks depend on the single drugs’ efficacy (*E*_1_,*E*_2_), and as a result, MSP frameworks are biased against the combination of moderately efficacious single agents. 3) The connection between the DEP and MuSyC is constrained by single drugs Hill slopes (*h*) and therefore DEP frameworks impose a Hill-slope dependent bias, artificially inflating the synergy for drugs with low Hill slopes. To assess impact of these considerations on synergy calculations in different fields, we analyzed two large publicly available datasets (Table 3) using MuSyC and other synergy frameworks (See Methods for description of fitting methods). Note, while synergistic cooperativity (*γ*) is theoretically plausible (as initially postulated by [9]), including it did not increase the fit quality (Supplemental Figure S3) as measured by AIC and therefore we ignore synergistic cooperativity in subsequent analysis.

**Table 3:** Summary of the datasets used for comparisons and validating theoretical predictions by MuSyC.

| Model | Number of Combinations | Metric of Drug Effect | Citation |
| --- | --- | --- | --- |
| *P. falciparum* (Strains: 3D7,HB3,Dd2) | 773 | Percent Response | [24] |
| 37 Cancer Cell Lines | 22738 | Percent Viable | [23] |

#### Box 1 Deriving Hill Equation from Mass Action Kinetics

Consider a reversible transition between an unaffected population (*U*) and an affected population (*A*) governed by

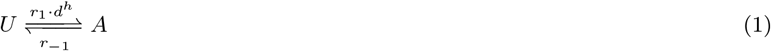

where *d* is the concentration of the drug, *h* is the Hill slope, often called cooperativity, and *r*_1_ and *r*_-1_ are constants corresponding to the reaction rate (Figure 1A). Applying the Law of Mass Action, steady state ratios of U and A are determined to be

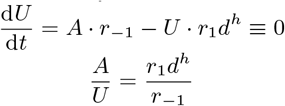

When 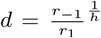, then (*A* = *U*). This dose is commonly called the EC50 (herein denoted as *C*). Because 100% of the population is either unaffected or affected, we also have the condition *U* + *A* = 1. This leads to the 2-parameter 1D Hill equation

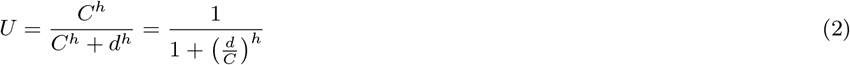

If the *U* and *A* differ by an observed effect (such as proliferation rate [26]), the measured effect *E* at dose *d* will be a weighted average

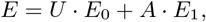

where *E*_0_ and *E*_1_ are the the effects characteristic of the *U* and *A*, respectively. From this we find the final form of a 4-parameter Hill equation:

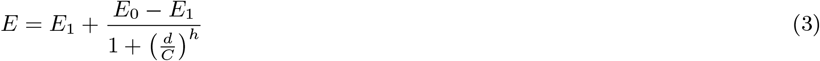

##### 2D extension of the Hill equation for two drug systems

Consider a system with 4 possible states, *U*, *A*_1_, *A*_2_, and *A*_1,2_ corresponding to populations that are unaffected, affected by drug 1 alone, affected by drug 2 alone, or affected by both drugs, respectively. The corresponding reactions between these states are:

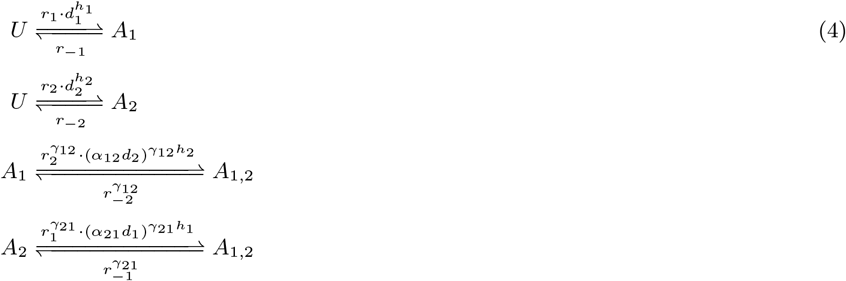

Here, the *α* parameters quantify the modulation of one drug’s EC50 (potency) due to the other drug. Similarly, the *γ* parameters measure the change of a drug’s Hill slope (cooperativity) due to the other drug.

As in the 1D case, finding the steady state of the system leads to the following system of equations

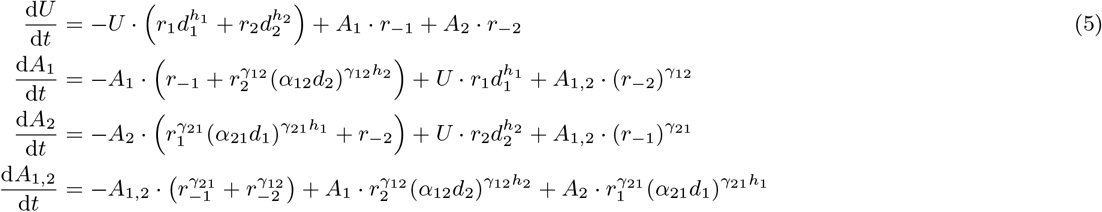

At equilibrium, the equations 5 must all be equal to zero; however, the system only defines a rank 3 matrix. Taking the first three equations from 5 with the constraint *U* + *A*_1_ + *A*_2_ + *A*_1,2_ = 1, we define

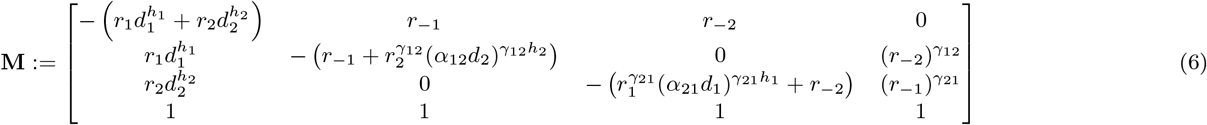

such that

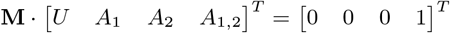

or, solving for the proportions of each state,

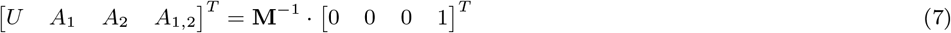

If we again consider distinct effects *E*_0_, *E*_1_, *E*_2_, and *E*_3_ distinguishing populations *U*, *A*_1_, *A*_2_, and *A*_1,2_, we find the equation for the dose response surface to be

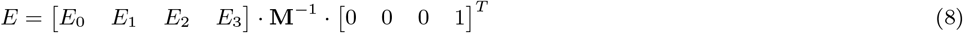

As *d*1 ← ∞ the equation reduces to

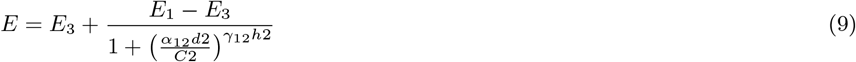

by which we can see the 2D equation reduces to a 1D Hill equation at the boundaries (See Supplemental Section *Proof of boundary behavior of 2D Hill equation*).

### Conflating synergistic potency and efficacy masks synergistic interactions

To determine how conflation of distinct synergy types impacts the interpretation of drug-response data, we generated synthetic dose-response surfaces using MuSyC (eq (8)) across a range of *α* and β values and calculated the synergy according to Loewe, Bliss, and Highest Single Agent (HSA) at the EC50 of both drugs (Figure 3A,D,G and Video S2). In each case, many distinct sets of (*α*_12_, *β*_21_, *β*) are indistinguishable (*e.g*. the black contour line on the spheres).

**Figure 3:**
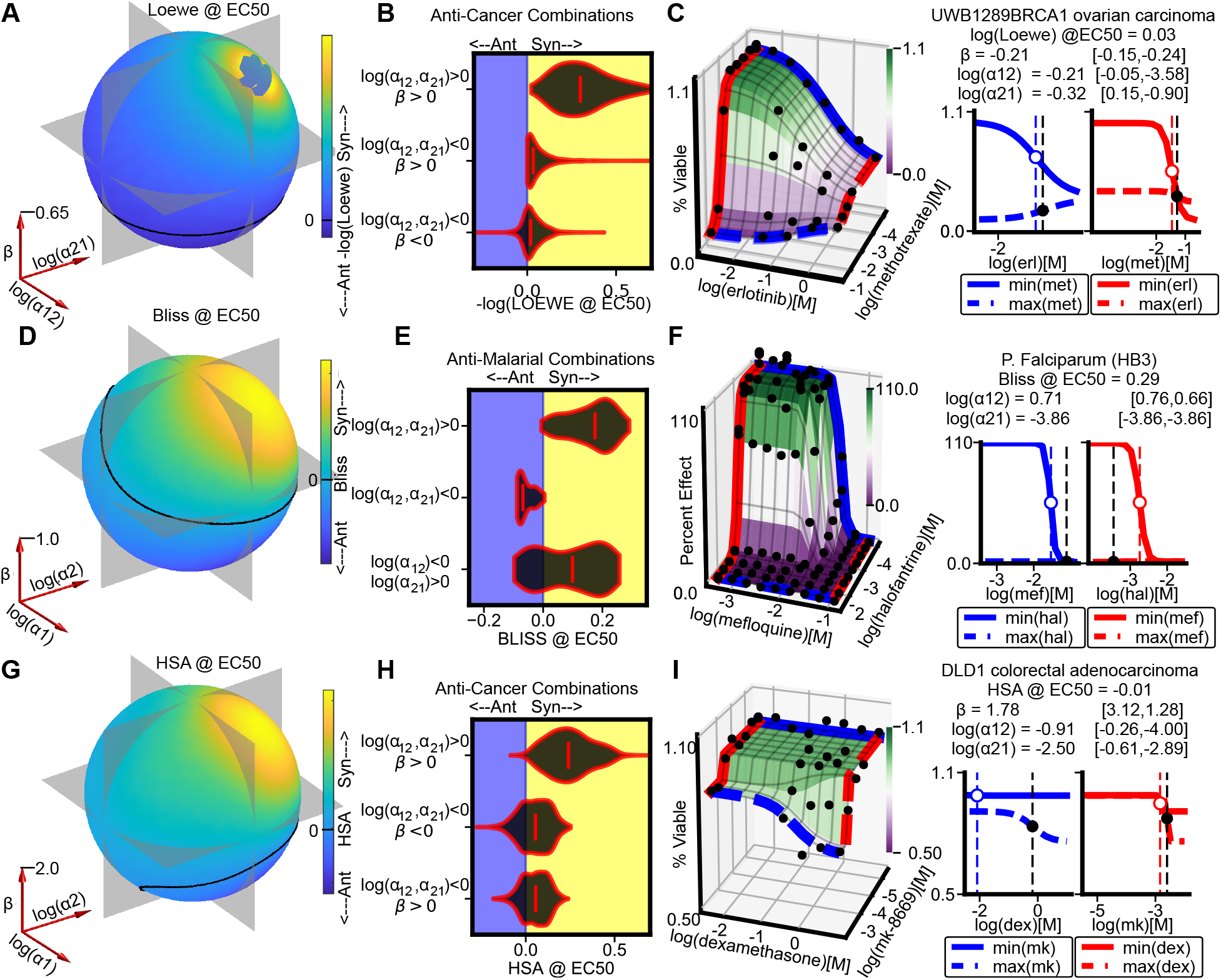
Conflating potency and efficacy synergy masks synergistic interactions in large drug combination datasets. A) The colors on the sphere (radius on *β* axis bottom left) represent the value of Loewe (colorbar to right) for a drug combination with a MuSyC synergy profile (*α*_12_, *α*_21_, and *β*) (axes bottom left). For all combinations: *E*_0_ = 1, *E*_1_ = *E*_2_ = 0, *h*_1_ = *h*_2_ = 1, *d*_1_ = *d*_2_ = *C*_1_ = *C*_2_, *γ*_12_ = *γ*_21_ = 1. The solid line marks the boundary between Loewe synergy and antagonism. Along this contour, which includes many different sets of (*α*_12_, *α*_21_, and *β*), Loewe is equivalent (-log(Loewe)= 0). Gray planes correspond to *β* = 0, log(*α*_12_) = 0, and log(*α*_21_) = 0. The hole in the upper-right quadrant represents sets for which Loewe is undefined. B) Distribution of Loewe for anti-cancer drug combinations grouped by their synergy profiles according to MuSyC. Loewe was calculated as detailed in Methods, including the Hill slope correction. C) The anti-cancer combination methotrexate and L-778123 is antagonistically potent and efficacious against HT29 cells, by MuSyC; however, it is designated by Loewe to be synergistic. Left panel shows the MuSyC-fitted dose-response surface, right panel shows the edges of the MuSyC surface. The open circle marks the EC50 for each drug in isolation, closed circle is the shifted EC50 due to antagonistic potency. Brackets are 95% confidence intervals for each parameter based on Monte Carlo sampling. D) Sphere for Bliss as in (A). E) Distribution of Bliss for anti-malarial drug combinations. Combinations for which each drug alone achieves *E_max_* < 0.1 were selected, ensuring *E*_1_ · *E*_2_ ≈ *E*_3_ ≈ 0. Under this condition, the differences between MuSyC and Bliss are due only to asymmetric potency synergy (all combinations fall near the *β* = 0 plane in (D)). F) Mefloquine increases the potency of halofantrine (red curves) but halofantrine decreases the potency of mefloquine (blue curves) in the HB3 strain of *P. falciparum*. G) Sphere for HSA as in (A). H) Distribution of HSA for anti-cancer combinations grouped by MuSyC synergy profile. In antagonistically potent combinations, HSA can miss synergistic efficacy. I) Combination of dexamethasone and mk-8669 in DLD1 cells is anagonistically potent, but synergistically efficacious.

Figure 3A shows that near the boundary between synergism and antagonism, Loewe is insensitive to changes in synergistic potency, tracking instead with synergistic efficacy. Consequently, in the anti-cancer dataset from O’Neil *et. al*. [23], Loewe misses potency antagonism in combinations with synergistic efficacy (Figure 3B middle distribution, see Figure S4 for an example surface). This reflects Loewe’s assertion of infinite potency antagonism (*α*_12_ = *α*_21_ = 0, Figure 2A) in its null model. Therefore, combinations that are antagonistically potent (*α* < 1) are all synergistic by Loewe in the absence of sufficient antagonistic efficacy (values above black contour in Figure 3A). Indeed, Loewe is frequently synergistic even in cases of antagonistic potency and efficacy (Figure 3B bottom distribution). As an example, the combination of methotrexate (targets folate synthesis) and erlotinib (EGFR inhibitor) in UWB1289 (BRCA1-mutant ovarian carcinoma) cells is antagonistically efficacious and potent by MuSyC, but synergistic by Loewe (Figure 3C).

Bliss synergy is classically thought to calculate synergistic potency. This is because assays where Bliss is appropriate (*E*_0_ = 1 and *E*_1_ = *E*_2_ = *E*_3_ = 0) by definition have no synergistic efficacy. However, even in these cases, Bliss still conflates *α*_12_ and *α*_21_ such that asymmetric potency synergy is obfuscated (Figure 3D, black contour line through *β* = 0 plane). In the anti-malarial dataset from Mott *et. al*. [24], Bliss is consistently synergistic when *log*(*α*_12_, *α*_21_) > 0, and antagonistic if *log*(*α*_12_, *α*_21_) < 0; however, when *log*(*α*_12_) < 0 < *log*(*α*_21_), Bliss will strictly classify a combination as either synergistic or antagonistic (Figure 3E bottom distribution) despite the asymmetric interactions. As an example, Bliss conceals that halofantrine (inhibits polymerization of heme molecules) reduces the potency of mefloquine (targets phospholipids) against the multi-drug resistant malaria strain HB3 (Figure 3F).

In contrast to Bliss, HSA is commonly thought to quantify synergistic efficacy. However, for antagonistically potent combinations, it cannot distinguish synergistic and antagonistic efficacy because it does not account for the topology of the dose-response surface (compare (log(*α*_12_), log(*α*_21_), *β*) = (−, −, +) and (−, −, −) quadrants of Figure 3G and Video S2). In the anti-cancer combination dataset [23], we observe this trend (Figure 3H middle vs bottom distributions). As an example, the synergistically efficacious combination of dexamethasone (agonist of the glucocorticoid receptor) and mk-8669 (PI3K/mTOR dual inhibitor) in a colorectal adenocarcinoma cell-line is masked by HSA due to antagonistic potency (Figure 1I). Repressing glucocorticoid signalling has previously been shown to repress mTOR signalling [27] providing a potential molecular mechanism explaining the synergy.

### MSP is biased against combinations of drugs with intermediate efficacy

MSP frameworks explicitly expect drug effects to measure the “percentage of cells affected,” which is by definition bounded within the closed interval *E* ∈ [0, 1]. Nevertheless, dose-response data is usually not a measure of percent *affect*, but rather of relative percent *effect* (see Supplemental Section *Percent Affect vs Percent Effect* for an example). This distinction, maintained by MuSyC (Box 1), is critical because percent effect data commonly saturates (*i.e*., percent *affect* is near 100%) at intermediate effect (*i.e*., relative percent *effect* is near 50%). For combinations of these moderately efficacious drugs, Bliss expects a large increase in effect over the single agents, even when each drug is administered at saturating concentrations (Figure 4A middle panel). In contrast, if combining drugs with high or low efficacy, Bliss expects a more modest increase (Figure 4A left and right panels).

Based on this expectation that *E*_3_ = *E*_1_ · *E*_2_ (Figure 2A), MuSyC predicts Bliss would be biased toward antagonism in combinations of moderately efficacious drugs (Figure 4B yellow shading around *E*_ı_ = *E*_2_ ≈ 0.5). As expected, the median Bliss score in the anti-cancer dataset is biased toward antagonism for moderately efficacious combinations 0.35 < (*E*_1_, *E*_2_) < 0.65 (Figure 4C, cyan square), though the magnitude of bias is less than predicted in Figure 4B. This bias persists even when looking at pan-cancer trends in the combination of drugs which have, on average, intermediate effect over the entire cell line panel (Figure 4D). As a particular example, the synergistic efficacy of paclitaxel (targets microtubule stability) and mk-2206 (AKT inhibitor) in KPL1 cells is masked by Bliss’s high expectation for moderately efficacious drugs (Figure 4E grey plane).

**Figure 4:**
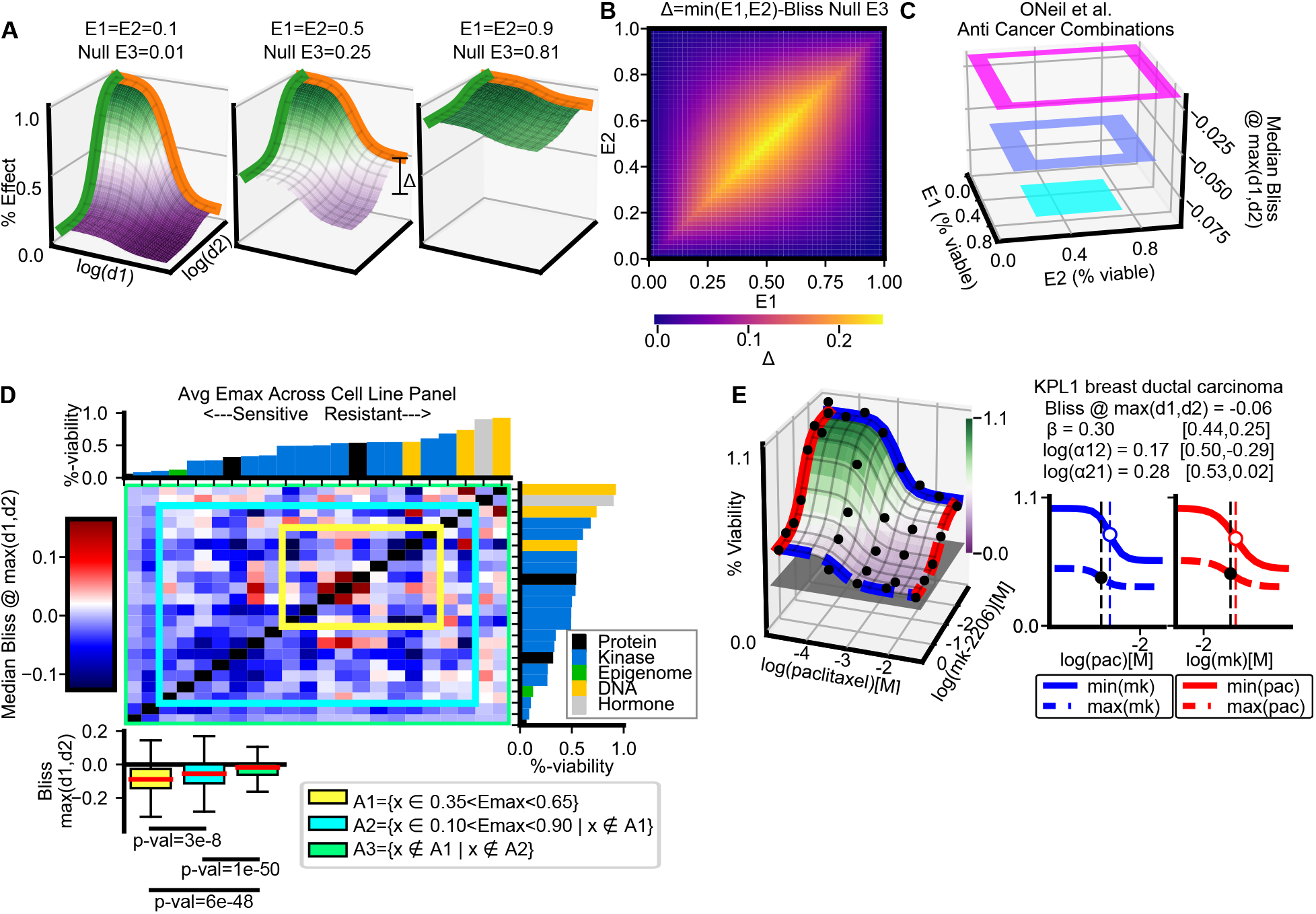
Bliss is biased against combinations of moderately efficacious drugs. A) The null Bliss surface for different maximal efficacy of single agents. Δ is defined as the expected increase in percent effect of the combination over the stronger single agent at saturating doses. The left and right panels have the same expected increase according to Bliss, Δ = 0.09, while the combination of moderately efficacious drugs (middle panel) has a expected increase of Δ = 0.25. B) Calculation of Δ (colorbar bottom) for surfaces with different pairings of (E1, E2). C) Median Bliss for anti-cancer combinations grouped by the maximal efficacy of their single agents. Ranges for each square: cyan square: [0.35, 0.65], blue square: [0.1, 0.9] and magenta square: [0.0, 1.0]. Bliss is calculated at the maximum tested concentrations of both drugs. D) Heatmap of the median Bliss score (colorbar left) for each combination across the cancer cell-line panel. Rows and columns are ordered by the average efficacy of each drug alone over all cell-lines (*E_max_*) (bar graphs top and right). Colored boxes correspond to groupings denoted in the legend (bottom). Boxplots show Bliss trends toward antagonism for combinations of moderately efficacious drugs (green->blue->yellow) (2-sample, 1-sided t-test). E) Dose-response surface of paclitaxel and mk-2206 in KPL1 cells. Gray plane is the expected effect of the combination by Bliss at max(d1,d2).

Additionally, some MSP methods, such as CI (nonexclusive) and the Effective Dose model, assume data measures percent *affect* and fit a simplified 2-parameter Hill equation enforcing *E*_0_ = 1 and *E*_1_ = 0. This assumption can lead to poor fits of percent *effect* data for moderately efficacious drugs, and thus invalid synergy scores (see Figure S5).

### Re-examining the sham experiment: Sham compliance introduces Hill-dependent bias in DEP models

A new synergy model’s consistency is traditionally tested with the “sham” combination thought experiment. In a sham experiment, a single drug is considered as though it were a combination, with the expectation that the drug should be neither synergistic nor antagonistic with itself. In Box 2, we show MuSyC only satisfies the sham experiment when *h* = 1. When *h* ≠ 1, the biochemistry of sham combinations (Figure 5A) is distinct from real combinations (Figure 5B), as states representing mixed-inhibition (black circles) are equivalent to single drug, complete-inhibition states (cyan circles) in sham combinations, but not in real combinations.

**Figure 5:**
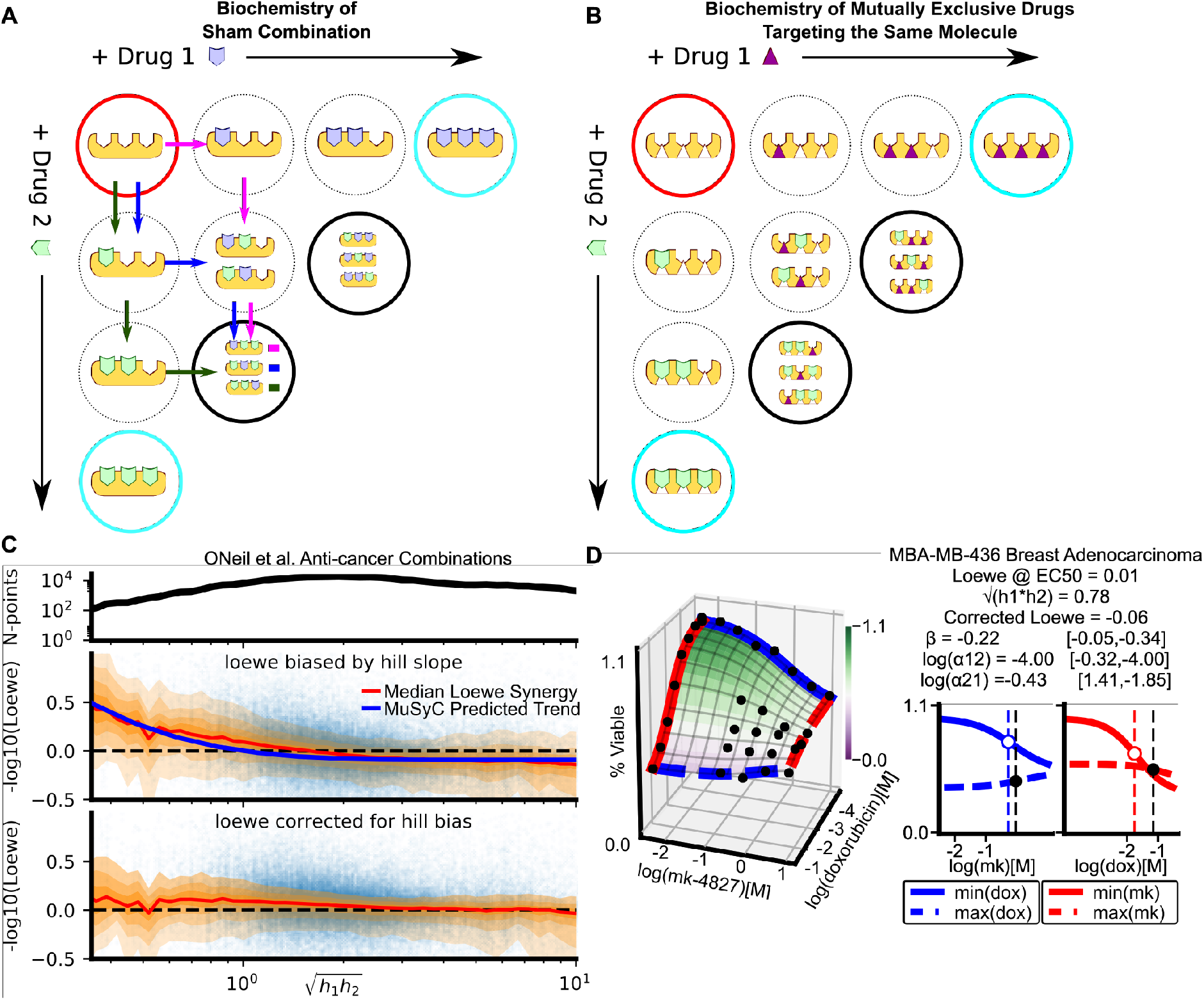
Enforcing sham compliance results in Hill-slope dependent bias in DEP frameworks. A) An illustration of the unique biochemistry of the sham experiment. The red circle represents an undrugged molecule with 3 binding sites. In a sham experiment, a drug is treated as though it were two separate drugs (green and blue polygons). Mixed states in which the binding sites are bound by both green and blue drugs (black circles) are equivalent to fully drugged states (cyan circles). We highlight three paths (green, blue, magenta arrows) that can be followed to reach a mixed-drugged state. These three paths correspond to the coefficient of 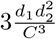 in equation (12) in Box 2. B) In a combination of mutually exclusive drugs (triangle and polygon), targeting the same molecule, and with the same number of binding sites, the mixed states (black circles) are not equivalent to fully drugged (cyan circles) accounting for the discrepancy between MuSyC and the sham experiment (Box 2). C) Loewe synergy is biased by Hill slope in the anti-cancer drug screen. The orange shaded regions show moving window percentiles (window width is 0.1) of Loewe (10th through 90th percentiles, in steps of 10). The top panel shows how many data points are present in the window. The blue curve in the middle plot shows the median MuSyC-predicted bias as a function of geometric mean of the Hill slopes (see Methods). Subtracting the MuSyC-estimated bias (calculated for each data point) from Loewe yields the bottom plot. C) The antagonistically efficacious and potent combination of mk-4827 and doxorubicin is misidentified as synergistic by Loewe, because both drugs in isolation have Hill slopes *h* < 1.

The constraint on h leads to non-linear isoboles in MuSyC (Figure S2) when *h* ≠1. Specifically, in the absence of synergistic potency (*α*_12_ = *α*_21_ = 1), MuSyC isoboles bend inward when *h* < 1, and outward when *h* > 1. Sham-compliant frameworks (Table 2) assume linear isoboles regardless of the Hill slope, and therefore classify combinations in the region between the linear and non-linear isoboles (red shading in Figure S2) as synergistic (middle panel) and antagonistic (right panel), respectively. MuSyC therefore predicts sham-compliant frameworks will overestimate synergy when *h* < 1, and underestimate when *h* > 1.

In combinations from the anti-cancer dataset, the average trend of Loewe synergy closely follows the Hill slope bias predicted by MuSyC (Figure 5C). Further, subtracting the MuSyC-predicted bias from Loewe values for each combination results in a distribution independent of Hill slope (bottom panel). The bias toward synergy is particularly large for drugs with low Hill slopes. As an example, both doxorubicin (DNA damaging agent) and mk-4827 (PARP inhibitor) have small Hill slopes when applied to MBA-MB-436 cells, and their combination is synergistic by Loewe. However, using MuSyC, we see this combination is both antagonistically efficacious and antagonistically potent (Figure 5D).

Therefore, satisfying sham compliance biases models toward synergy for drugs with low Hill slopes, regardless of with what these drugs are combined. This bias—which stems from enforcing a biochemical reaction scheme only appropriate for sham combinations—should be sufficient grounds for dismissing the sham experiment as a measure of a new synergy framework’s validity.

#### Box 2 Sham compliance of MuSyC, and the mass action biochemistry of a sham experiment

To simulate a sham experiment using MuSyC, there is no state *A*_1,2_ (Figure 1B), which requires *α*_12_ = *α*_21_ = 0. Further, because drugs 1 and 2 are the same, *h*_1_ = *h*_2_ = *h*, *C*_1_ = *C*_2_ = *C*, and *E*_1_ = *E*_2_. Thus, the 2D Hill equation (eq. (8)) reduces to

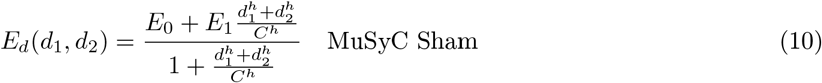

In comparison, the true dose-response surface of a sham experiment can be analytically determined from the 1D Hill dose-response equation (eq. 3) as

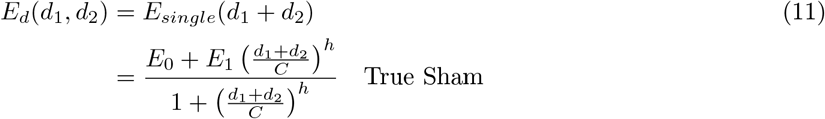

Equations 10 and 11 are only equivalent when *h* = 1. This makes sense, as the constraints on *α* and *h* are the conditions required for MuSyC to satisfy the DEP (Figure 2B). To see what happens when *h* ≠1, consider, for instance, a molecule with three binding sites targeted by a small molecule inhibitor (*h* = 3). For clarity, we assert *E*_0_ = 1 and *E*_1_ = 0, though the findings are valid more generally. The MuSyC sham surface follows

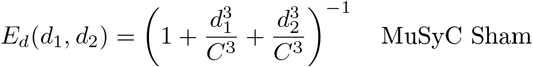

In contrast, the true sham surface is

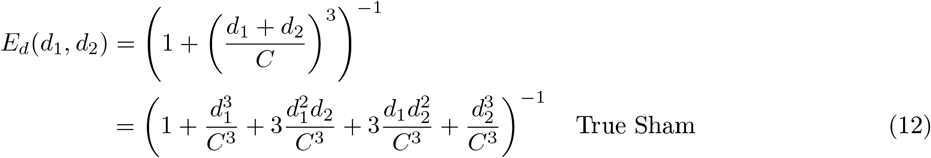

The two additional cross-terms in the true sham equation (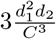 and 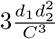) describe the six possible mixtures of drugs 1 and 2 that, together, fill all binding sites (Figure 5A, blue, green, and magenta paths show three possible mixtures). In a sham experiment, because drugs 1 and 2 are the same, the diagonal states (black and cyan circles) in Figure 5A are all equivalent, and fully inhibited.

Conversely, in non-sham combinations, drugs rarely target the same binding sites, or even the same molecule. Even when two drugs are mutually exclusive inhibitors of the same molecule *and* have the same number of binding sites, the cross-terms describe non-equivalent, not fully inhibited states (Figure 5B). A commonly applied and physiologically supported approximation is that only fully bound molecules become (in)active (see reaction schemes 5-7 in [28]). Partially bound cross-terms are therefore reasonably modeled as unaffected, and the absence of these cross-terms from equation (10) is justified for real (non-sham) combinations (see Discussion). Further, when the two drugs do not target the same molecule or are mutually exclusive or have the same number of binding sites, by far the preponderance of real combinations, the diagonal states are ill defined yet remain embedded in the sham equation.

## Discussion

Herein, we have demonstrated three key advances of MuSyC [21] particularly germane to the study of combination pharmacology: 1) the unification of the DEP and MSP; 2) the decoupling of three distinct types of synergy; and 3) the revelation of biases emerging from constraints on the single drug pharmacological profile inherent in the DEP and MSP.

The DEP and MSP have formed the foundational principles of most synergy frameworks over the last century; however, the connection between these principles has remained unknown [6, 7]. Here, approaching combination pharmacology using the Law of Mass Action results in a single framework unifying both principles. By mapping all frameworks on a common landscape, MuSyC facilitates rigorous investigation of oft-cited, contradictory conclusions between existing frameworks [5]—contradictions that preclude reproducibility between synergy studies. Specifically, as is seen in Figure 2C, there is no combination which can simultaneously satisfy the conditions required by both DEP and MSP synergy frameworks.

One key advance of MuSyC, facilitating this unification, was the decoupling of *α*, *β*, and *γ*. These synergy parameters correspond directly to classic, pharmacological measures of a drug’s potency, efficacy, and cooperativity. By calculating synergy in this way, its interpretation does not depend on arbitrary expectations or thresholds. Rather, an *α* of 10 corresponds to a 10-fold increase in a compound’s potency, as a result of the other drug, regardless of whether we define *α* = 1 or *α* = 10 as the “threshold” for synergy. We envision distinguishing synergies of potency, efficacy, and cooperativity will be of differential consequence in different contexts. For example, in cancer synergistic efficacy may be most important, while for neurological disorders, synergistic cooperativity —to increase the switch-like behavior of drugs —may be preferred.

The relationship between MuSyC and the MSP and DEP frameworks (Figure 2) is constrained by single-drug parameters (*E*_1_,*E*_2_ for MSP, *h* for DEP). These constraints suggested systematic biases in MSP and DEP frameworks contingent on a single drug’s efficacy (MSP) and Hill slope (DEP). These systematic biases merit consideration when using these frameworks for drug discovery in large screens. Additionally, the constraint on h highlighted a discrepancy between the biochemistry of true sham experiments and real combinations. The centrality of the sham experiment in the drug synergy literature cannot be overstated; however, we argue enforcing sham compliance comes at the cost of improperly modeling real combinations, leading to a predictable Hill-dependent bias.

The prospects of higher order synergies (*i.e*., interactions beyond pairwise) and scaling laws for drug mixtures, while provocative, have remained contentious [29, 8, 30]. MuSyC’s cubic geometry allows it to be easily extended to three or more drugs (Figure S1), and we expect MuSyC will enable a more refined search for higher order interactions. For instance, combinations that mix different synergy profiles (*e.g*., drugs 1 and 2 are synergistically potent, and drugs 2 and 3 are synergistically efficacious) may exhibit different higher order interactions than combinations all sharing a single synergy type. However, the number of synergy parameters in MuSyC scales poorly, and the commensurate data necessary to fully constrain MuSyC hyper-surfaces invokes a parameter identifiability problem. MuSyC’s geometry could be leveraged to guide sampling schemes to constrain the boundaries, allowing the solution to be built up step-wise (see Supplemental Section *Proof of boundary behavior of the 2D Hill equation*).

MuSyC expects single-drug dose-response curves to be sigmoids well fit by the 1D Hill equation (eq (3)), and dose-response surfaces to be well fit by the 2D Hill equation (eq (8)). In our experience, these expectations are met by real data, as most single drugs have monotonic, sigmoidal responses, and even complex drug interactions can be modeled using various mixtures of *α*, *β*, and *γ* (96% and 88% of combinations in anti-cancer and anti-malarial datasets had *R*^2^ > 0.7, respectively). However, it is possible for drugs to have multiphasic responses due to poly-pharmacology which is not well fit by a Hill curve. It may be possible to extend MuSyC to encompass such drugs —for instance by including a multiphasic Hill model [31] or modeling effects of “partially affected” states (Figure 5A,B and Box 2). In extreme cases, it may only be possible to apply non-parametric frameworks such as Bliss, Loewe, or HSA. Additionally, MuSyC assumes all drugs are administered concurrently, whereas patient treatments are often staggered. New theory and experimental methods are needed to address the synergy of combinations which are staggered temporally, bridging the synergy of pharmacodynamics with the synergy of pharmacokinetics. Finally, in the datasets we analyzed here, we did not find a role for *γ*. Future studies in other systems are needed to better understand situations when synergistic cooperativity is expected.

By viewing the landscape of drug synergy through the lens of mass-action, we have demonstrated the underlying assumptions, limitations, and biases of commonly applied synergy methods. We have shown how MuSyC unifies the DEP and MSP thus providing a consensus framework for the study of combination pharmacology. These findings provide much needed clarity to the field and should promote the reproducibility of drug synergy studies across drug combination discovery efforts. Such a rigorous approach to the discovery and application of drug combinations will open the door to the discovery of new and previously discarded avenues for therapeutic mixtures.

## Methods

We note the analyses conducted were not necessarily the same as those used in the original paper. Indeed the limitations of the current frameworks forced customized analysis for each publication highlighting the need for a consensus framework. Here we describe fitting protocol for drug metrics where the metric of drug effect *decreases* as dose increases (*E*_0_ > *E*_3_) (*e.g*., anti-proliferative drugs); however, the framework is equally valid if increasing the drug corresponds to increases the effect (*E*_0_ < *E*_3_) (*e.g*., percent effect)

### Fitting 2D Hill equation

The following packages were used for fitting, data analysis, or visualization: GNU parallel [32], SciPy [33], Numpy [34], Pandas [35], Matplotlib [36], uncertainties [37]. Fitting was done using a custom library written in Python2. Previously, we found it necessary to use a Metropolis Hastings Monte Carlo (MCMC) seeded with a particle swarm optimization (PSO) to fit the 2D Hill equation [21]. This was prompted by the inconsistent performance of standard non-linear least squares (NLLS) regression. In particular, we observed instances of calculated uncertainties in NLLS which were several orders of magnitude greater than the parameter value. This, we have discovered, is because the multi-collinearity between the Hill slope and the EC50 (*C*) inherent in the structure of the Hill equation—collinearities which are amplified when extending the Hill equation to 2D. The correlation between variable *h* and *C* is easiest to observe in a linearized 1D Hill equation (eq 13).

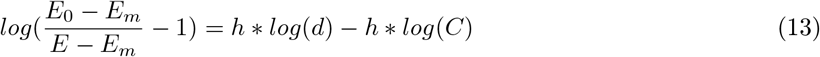

In eq. 13, the intercept of the line (*h* * *log*(*C*)) depends on the slope of the line (*h*). This correlation is problematic when trying to estimate the parameter uncertainty (*σ*) from NLLS regression because *σ* is estimated as the square root of the inverse Hessian, approximated to be *J^T^ J* (where *J* is the Jacobian at the solution). *J* of the 4 parameter Hill equation is

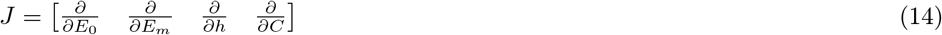

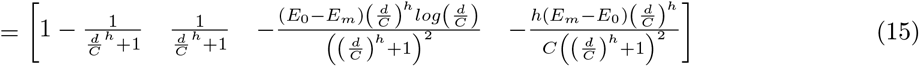

When the Hill slope is large (*e.g*., h>5), the second two terms of the *J* cause the inverse of the Hessian matrix to be undefined. This problem is compounded in the 2D Hill equation where, in addition to *h* and *C*, the parameters *α* and *γ* are co-linear. However, this does not affect the accuracy of the fitted parameter values from the NLLS regression—only the parameter uncertainty [38].

For the fitting the 2D Hill equation in this study, we adopted a Monte Carlo sampling approach to calculate the fit parameter uncertainty. This is significantly faster than our previous method (PSO+MCMC) [21] while maintaining reasonable calculations of the parameter uncertainties accounting for the multi-collinearities described above. The Monte Carlo algorithm for fitting the 2D Hill equation is as follows. First, the 4-parameter 1D Hill equation (eq. 3) is fit to the dose-response of each drug in isolation. The fit uses the Trust Region Reflective (TRF) algorithm implemented in the *curve_fit*() module of the scipy.optimization package. *h* and *C* were unconstrained while *E_max_* and *E*_0_ are constrained for each dataset as annotated in the methods section *Data acquisition, preparation, and analysis*. The initial 1D Hill fits provide estimates for (*E*_0_, *E*_1_, *E*_2_, *C*_1_, *C*_2_, *h*_1_, *h*_2_). Next the 2D Hill equation (eq. 8) is fit using the TRF algorithm with initial values based on the 1D Hill equation fits and with bounds based on the parameter uncertainty calculated for the 1D Hill fits. The initial values for parameters unique to the 2D Hill equation, *E*_3_, *α*_21_, *α*_12_, *γ*_12_, *γ*_21_ are (*min*(*E*_1_, *E*_2_),1,1,1,1). For all combinations *r*_1_ = *r*_2_ = 100. The bounds for log(*α*_21_), log(*α*_12_) are set to [−4,4]. From this initial fit, 100 Monte Carlo samples are used to calculate the parameter uncertainty as described by Motulsky and Christopoulos [38], (*Chapter 17: Generating confidence intervals by Monte Carlo, pg. 104*). Specifically, noise, with a distribution N(0,*σ*), where *σ* is equal to the root mean square (RMS) of the best fit, is added to best-fit values of the 2D Hill equation for all drug doses. The data plus noise is then fit again initializing the optimization from the best fit parameters of the original data. This is done 100 times. From this ensemble of fits, the 95% confidence interval of each parameter can be calculated. This Monte Carlo approach results in asymmetric confidence intervals which better captures the non-Gaussian distribution of uncertainty for many fits (*e.g*. the distribution of *h* is log-normal) as well as being robust to the co-linear parameters in the 2D Hill equation. The asymmetric confidence interval is particularly salient when the dose-range is insufficient to observe the lower plateau of the dose-response. Only combinations for which *R*^2^ > 0.7 and the fitted EC50s of both drugs was less than maximum tested dose for each (*C*_1_ < *max*(*d*1),*C*_2_< *max*(*d*2)) were included for subsequent analysis.

### Data acquisition, preparation, and analysis

#### ONeil et al. Anti-cancer Screen

The anti-cancer drug combination data was downloaded from the supplemental materials of [23]. Single agent and combination datasets were merged. Drug effect was the mean normalized percent viability (*X/X*_0_ column) calculated as detailed in [23]. The minimum and maximum bounds for [*E*_0_, *E*_1_, *E*_2_, *E*_3_] during 2D Hill equation fits were [.99,0.,0.,0.,0.] and [1.01,2.5,2.5,2.5] respectively.

#### Mott et al. Anti-malarial Screen

The anti-malarial drug combination data was downloaded from https://tripod.nih.gov/matrix-client/ from the **Malaria Matrix** project. Blocks downloaded for analysis were: [1601,1602,1603,1701,1702,1703,1761,1764]. Only blocks with a *mqcConfidence* of greater than 0.9 were included. The drug effect was calculated as described in [24]. Effects less than −20% and greater than 120% were removed. The minimum and maximum bounds for [*E*_0_, *E*_1_, *E*_2_, *E*_3_] during 2D Hill equation fits were [90.,0.,0.,0.] and [110,200,200,200] respectively.

### Calculation of other synergy metrics

#### Bliss, Loewe, and HSA

Bliss, Loewe, and HSA depend on the concentration of drugs so a combination can be synergistic at one dose, but antagonistic at another dose. Several methods have been proposed for extracting a single synergy metric per combination from these frameworks to enable comparisons between combinations [14, 23, 15, 16]. For our analysis, we calculate the synergy score at the combination of each drug’s EC50 (Figure 3,5) or at the maximum tested concentration of each drug (Figure 4). The EC50 of each drug was calculated from the fits to the 2D Hill equation ((8)) which we have observed to be more robust to noise when estimating the single drug pharmacologic profile. Assuming the notation for the 1D Hill equation and inverse Hill equation—which calculate effect (*E* given a dose (*d*) and a dose given an effect, respectively—are given by

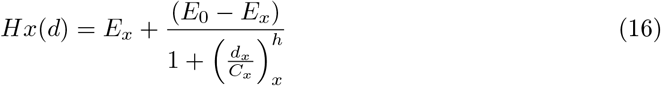

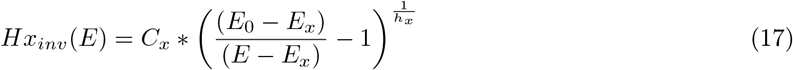

where *E_x_* < *E*_0_, then equations for Bliss, Loewe, and HSA at the EC50 are:

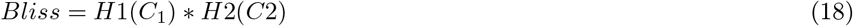

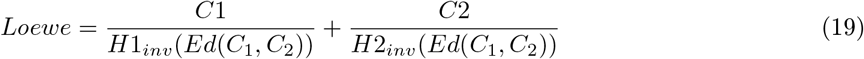

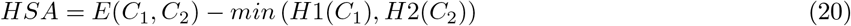

where *Ed*(*C*_1_, *C*_2_) is the measured effect of combining *C*_1_ of *d*_1_ and *C*_2_ of *d*_2_. And equations for Bliss at the max of each drug is:

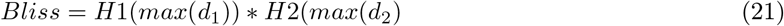

These equations assume the metric of drug effect decreases as the dose increases. Because many single agents did not reach 0% maximum efficacy, the EC50s (*C*_1_, *C*_2_) were not necessarily 50% (Figure S5). If *E*(*C*_1_, *C*_2_) < *E*_1_, *E*_2_ then Loewe was undefined. We apply a − log_10_ transformation the scale Loewe to match the ranges Bliss and HSA are synergistic; therefore, f − log_10_ (*Loewe*) > 0 the combination is synergistic and if − log_10_(*Loewe*) < 0 the combination is antagonistic. Additionally, for Figure 3 and Figure 5 we had to calculate the Hill-dependent bias in Loewe. For Figure 3, we subtracted the Hill slope bias to only study the impact of conflating synergistic potency and efficacy. To calculate the bias, Loewe was calculated as in equation (19) where was was evaluated at the MuSyC-predicted *Ed*(*d*_1_, *d*_2_) for the combination retaining all the single drug parameters (*E*_0_, *E*_1_, *E*_2_, *C*_1_, *C*_2_, *h*1, *h*2) and assuming (*α*_12_ = *α*_21_ = 0). This resulted in an estimate of the bias purely due to the Hill slope in the Loewe calculation.

#### ZIP and BRAID

Both ZIP and BRAID were calculated for each combination using the R packages available for each method:

(ZIP’s R code is in the supplemental file 1 of the manuscript [9] and BRAID’s package is available from: https://cran.r-project.org/web/packages/braidReports/braidReports.pdf).

#### Effective Dose Model

To fit Zimmer et al.’s Effective Dose Model we used the `scipy.optimization.curve_fit` module in Python 2.7. Specifically, the Effective Dose Model, equation 2 in [39] (eq. (30) in Supplement), contains parameters (*C*_1_, *C*_2_, *a*_12_, *a*_21_, *h*_1_, *h*_2_) where the a parameters are the synergy values. In the model, there are no parameters for efficacy because it is assumed the drug effect ranges between zero and one. When this is not true, the Effective Dose Model results in poor fits to the data (Figure S5).

#### Schindler’s Hill PDE model

The Hill PDE model has no parameters to fit as the dose-response surface is derived the single dose-response curves. In fact, Schindler does not propose a method to estimate synergy from experimental data, but postulates some implementation of perturbation theory could be used to fit experimental data [10]. Therefore, to calculate the synergy of this model, we defined the sum of residuals between the null surface and the experimental data to the metric of synergy.

#### Combination Index (CI)

Following the CI fitting algorithm in the CompuSyn software, we fit the median-effect equation, a 2-parameter, log-linearized form of the Hill equation obtained by assuming *E*_0_ = 1 and *E*_1_ =0 [3]. *C* and *h* were then obtained from the slope and y-intercept of the log-linearized data using the `scipy.stats.linregress` module in Python 2.7. While CI assumes the drug effect is scaled between (0,1), when this is not the case, a poor fit results (Figure S5A). All data points with percent viability greater than 1 were excluded from the CI fit because the median-effect equation becomes complex in those cases. For some drugs, this left too few points to fit a line, such that CI was undefined. In other cases, the fitted value for h was less than zero, a physically unrealistic result; therefore, those combinations were also considered undefined. After that, CI was calculated as

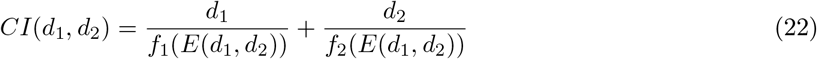

where *f_i_*(*E*) is obtained by solving the Hill equation for *d*, and is given by

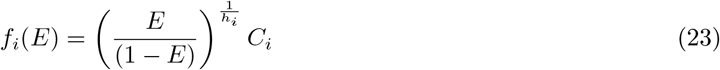

As with Loewe, we apply a −log_10_ transformation to scale CI synergy such that −log_10_(*CI*) > 0 the combination is synergistic and if −log_10_(*CI*) < 0 the combination is antagonistic.

## Videos

**Video S1:** Synergistic efficacy (*β*), synergistic potency (*α*), and synergistic cooperativity (*γ*) correspond to orthogonal geometric transformations of the dose response surface.

**Video S2:** Calculations of Loewe, Bliss and HSA for different sets of (*α*_12_, *α*_21_, *β*) on a spherical manifold. Color represents value of Loewe (left), Bliss (middle), and HSA (right). Radius and orientation of the sphere is depicted on bottom left axis. Where Loewe is undefined the sphere is blank.

## Acknowledgments

The authors would like to thank Corey Hayford, Sarah Maddox, Darren Tyson, Leonard Harris, Joshua Bauer, James Pino, Ken Lau, and Chris Wright for insightful conversations and critical feedback. This work was supported by the following funding sources: CTM was supported by National Science Foundation (NSF) Graduate Student Fellowship Program (GRFP) [Award #1445197]; CFL was supported by the National Science Foundation [MCB 1411482]; CFL and VQ Were supported by the National Institutes of Health (NIH) [U54-CA217450 and U01-CA215845]; VQ was supported by NIH [R01-186193].

## Author Contributions

All authors conceived and designed the analyses and reviewed the final manuscript; CTM and DJW performed the analyses and visualizations; CFL, CTM, and DJW wrote the manuscript.

## Supplemental Materials

### 1 Supplement Figures

**Figure S1:**
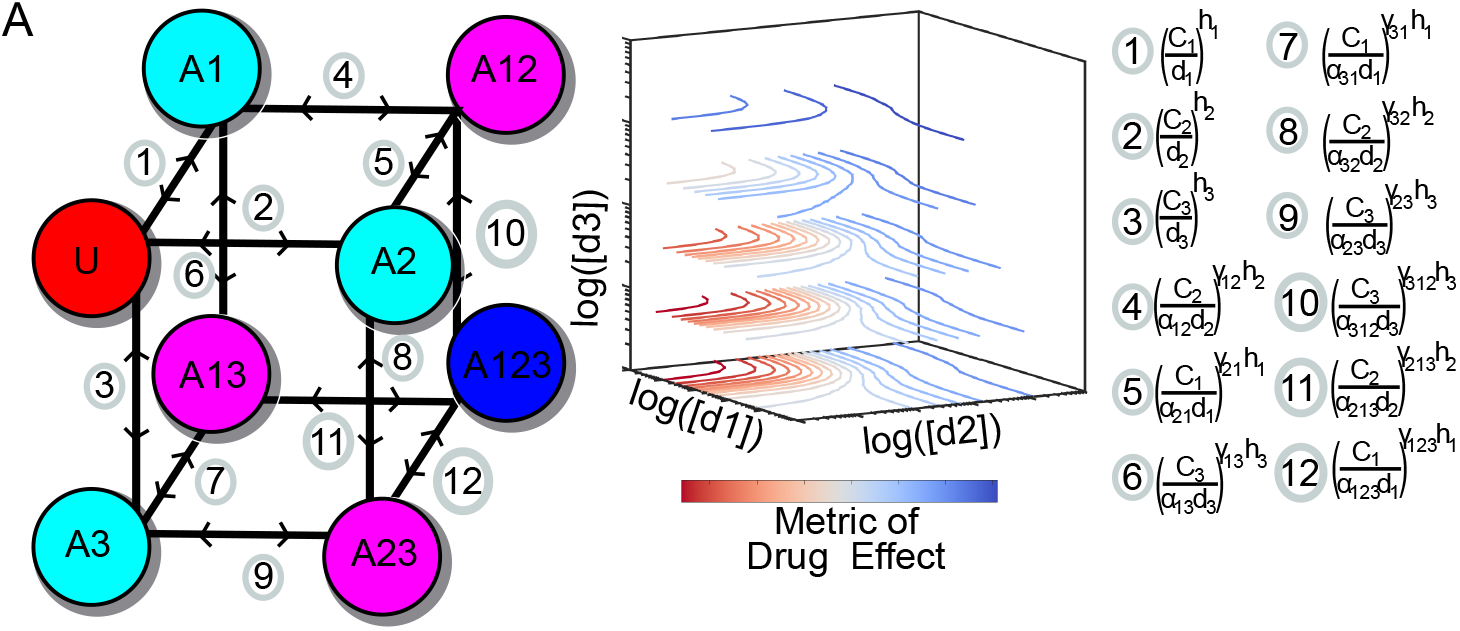
A) Extension of MuSyC to combinations of three drugs. Following the cubic geometry of Figure 1B, combinations of 3 drugs result in a cube. Numbered notation next to each edge corresponds to the ratio of the connected corners at equilibrium for the boundary conditions. For example, edge #10 annotation means 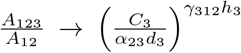 as *d*_1_ → inf, *d*_2_ → inf. For combinations of 4 drugs, the geometry is a tesseract. In general, MuSyC can describe combinations *N*-drug combinations by considering 2^*N*^ possible states with transitions defining the edges of an *N*-dimensional hypercube. Dose-response surfaces generalize to *N*-dimensional scalar functions. In the most general case, for *N* drugs there are 2^*N*^ − *N* − 1 distinct *β* parameters (one for each state characterized by the action of at least 2 drugs), and *n* · (2^*n*−1^ − 1) distinct *α* and *γ* parameters (one for each edge, excluding edges connected to the undrugged state, which correspond only to single-drug potency and cooperativity). Thus, MuSyC can account for higher-order synergyies (*e.g*., synergy that emerges from a combination of three drugs, but is not evident in any pairwise combination of those drugs), however the rapid growth of the number of synergy parameters with N suggests that significant quantities of data, or confident knowledge of pairwise synergies, would be needed to measure such higher-order synergies.

**Figure S2:**
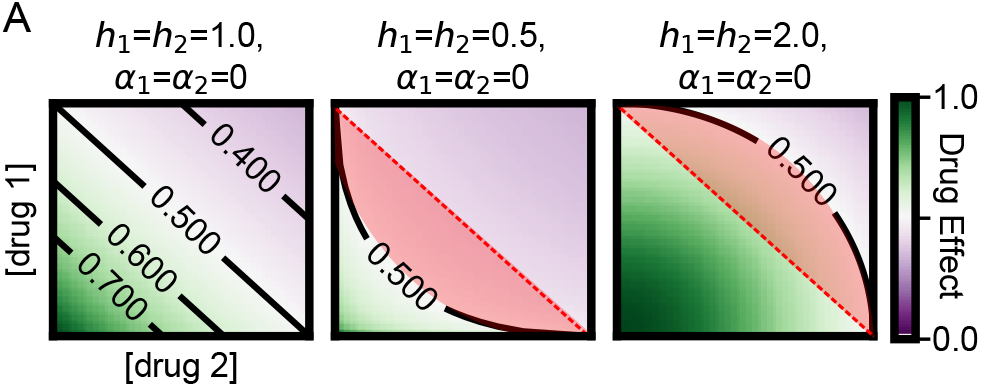
A) When *h* =1 and *α* = 0, MuSyC results in linear isoboles (contours of equivalent effect) characteristic of DEP models. When *h* < 1 (middle panel), the isoboles bend inward such that DEP models will misclassify the red region as synergistic, biasing DEP calculations toward synergy for *h* < 1. Conversely when *h* > 1, isoboles bend outward (right panel) and DEP models will misclassify the red region as antagonistic, biasing Loewe toward antagonism for *h* > 1.

**Figure S3:**
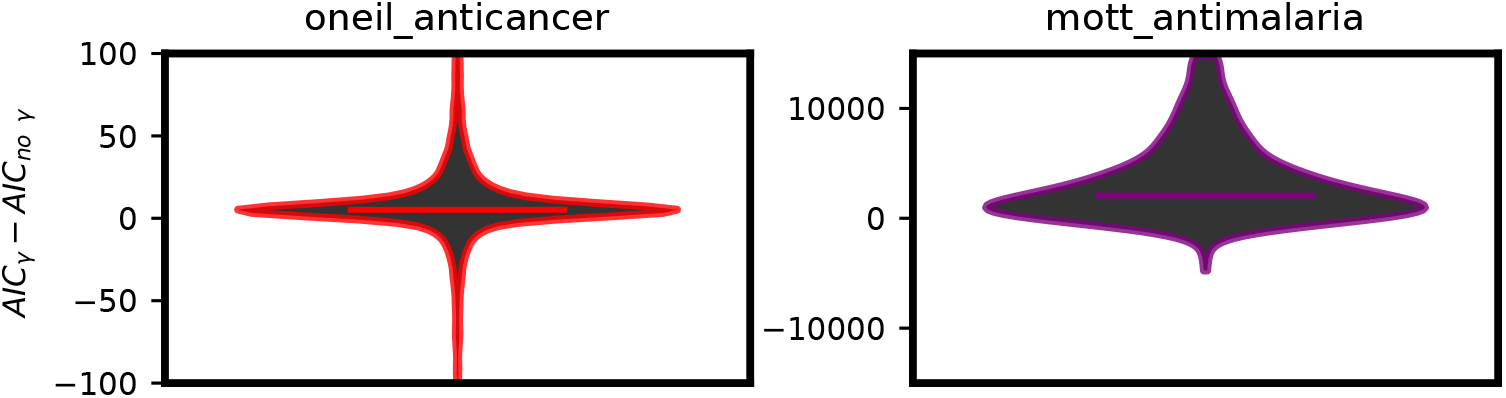
A) Difference in the Akaike Information Criterion (AIC) values for MuSyC models including fitting synergistic cooperativity (*γ*_12_ and *γ*_21_) or fixing *γ* to 1 thereby reducing the parameter count by 2. Models which minimize AIC are preferred. In most cases the simpler model is preferred. The mean *AIC_γ_* − *AIC_noγ_* for anti-cancer and anti-malarial datasets was 8 and 3312, respectively. The percent of combinations for which the model including *γ* had a lower AIC value was 18% and 5%, respectively.

**Figure S4:**
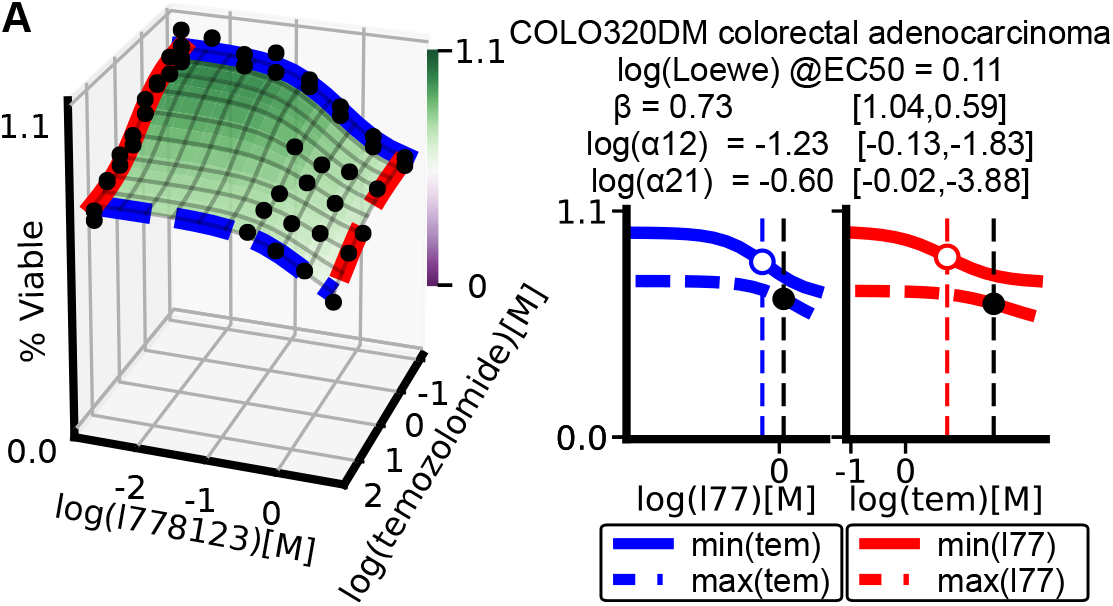
A) The combination of L778123 (a dual farnesyl and geranylgeranyl transferase inhibitor) and temozolomide (DNA alkylating agent) in COLO320DM cell lines is synergistically efficacious but antagonistically potent, and is an example where Loewe misses antagonistically potent interactions.

**Figure S5:**
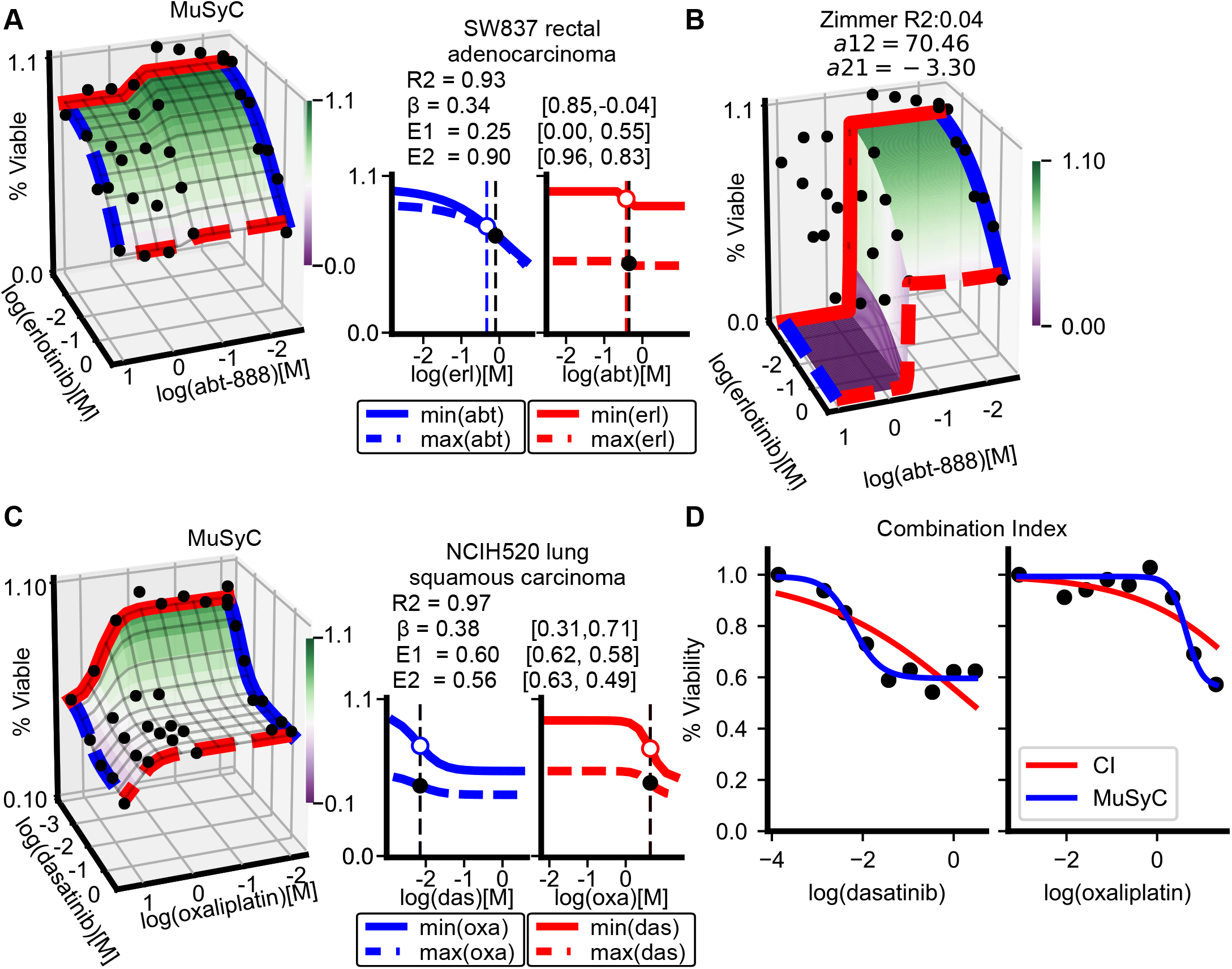
A) Dose-response surface for combination erlotinib and abt-888 in SW837 cells. B) Dose response surface according to Effective Dose model, which enforces *E*_0_ = 1 and *E*_1_ = *E*_2_ = 0. C) Dose-response surface for dasatinib and oxaliplatin in NCI-H520 cells. Both dasatinib and oxaliplatin have intermediate effects on this cell line (*i.e. E*_1_ ≈ *E*_2_ ≈ 0.5). D) CI single drug dose-response fits for dasatinib and oxaliplatin. Minimizing residuals while enforcing *E*_0_ = 1 and *E*_1_ = *E*_2_ = 0 causes artificially low Hill slopes for both drugs (*h*_1_ < *h*_2_ < 0.35). For additional examples of poor CI fits see [21].

**Figure S6:**
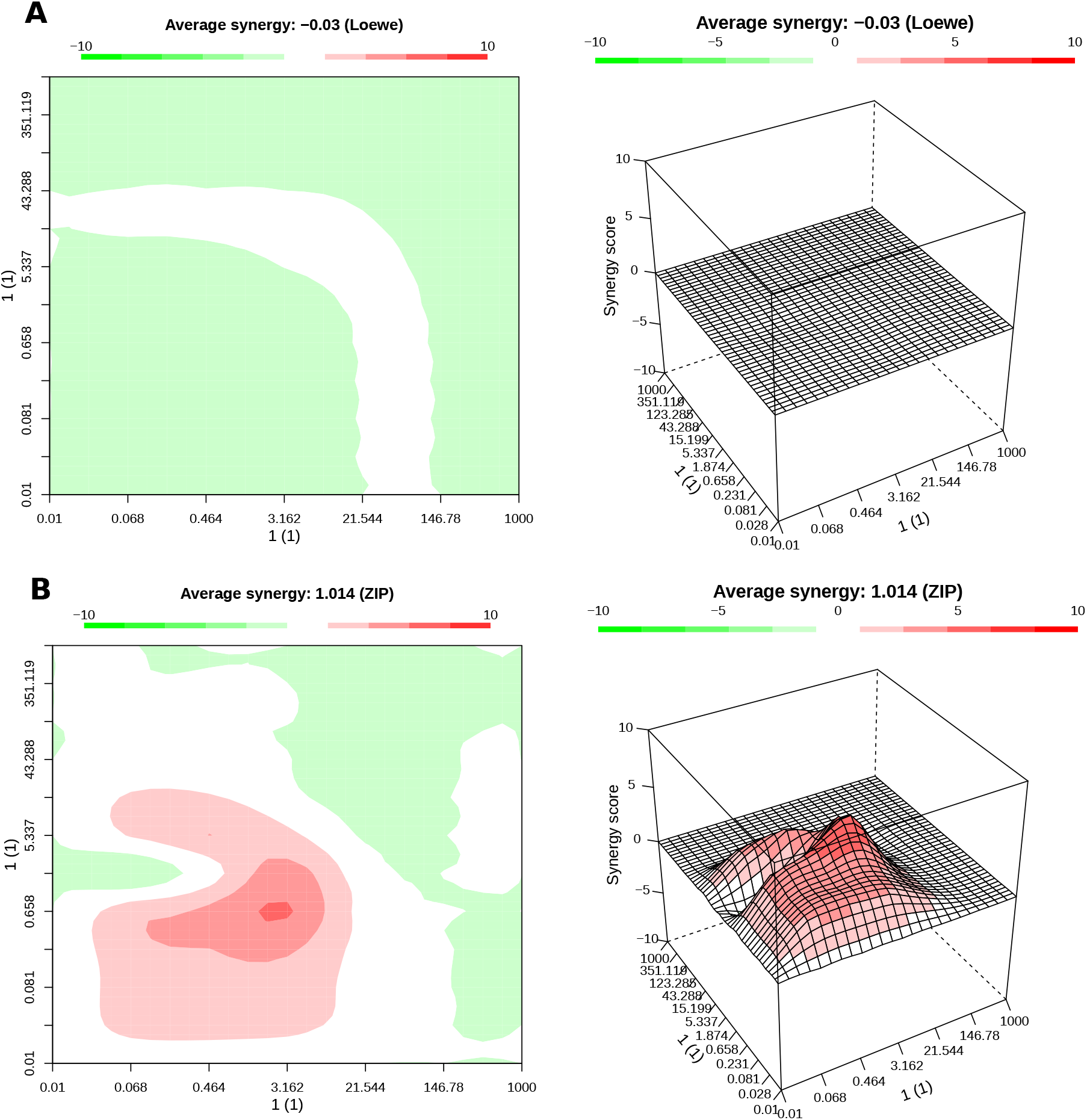
A) Loewe synergy calculated by synergyfinder [16] for a synthetic sham dose-response surface with *h* = 2. Loewe correctly identifies the combination as additive. B) ZIP quantifies synergy or antagonism at several concentrations for the sham dataset.

### 2 Relationships between MuSyC and the MSP and DEP

#### 2.1 Multiplicative Survival Principle

MuSyC matches the Bliss null surface when there is no potency synergy (*α*_12_ = *α*_21_ = 1), no cooperativity synergy (*γ*_12_ = *γ*_21_ = 1), and *E*_3_ = *E*_1_ · *E*_2_ (Figure 2A). To show this, let each drug in isolation have a 1D hill response

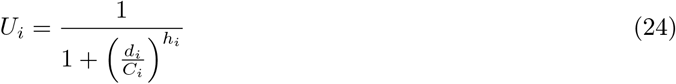

where *U_i_* reflects the portion of cells unaffected by drug *i* alone. For the 2D case, when *α*_12_ = *α*_21_ = 1 and *γ*_12_ = *γ*_21_ = 1, each edge in Figure 1B satisfies detailed balance and therefore the state occupancy is given by

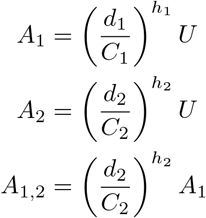

Because *U* + *A*_1_ + *A*_2_ + *A*_1,2_ = 1, the MuSyC mass action model gives

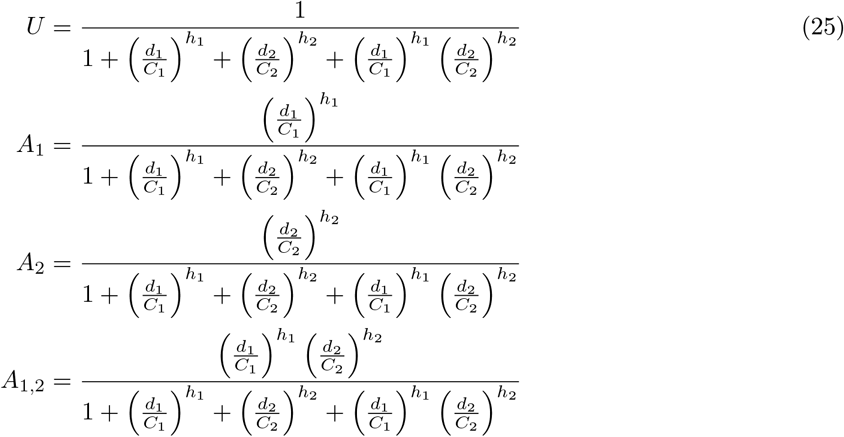

From this, it is easy to verify that *U* = *U*_1_ · *U*_2_ where *U*_1_ = 1 − (*A*_1_ + *A*_1,2_) and *U*_2_ = 1 − (*A*_2_ + *A*_1,2_ which is equivalent to the Bliss Independence null model.

Furthermore, given *E*_0_ = 1

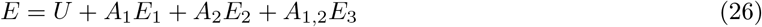

We define 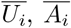, and 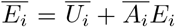 to be the fraction of unaffected cells, fraction of affected cells, and observed effect for treatment due to the single drug *i*, as described by equation 24. The overline distinguishes affects attributable to each drug, such that 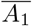 includes cells affected either by drug 1 alone, or by both drug 1 and drug 2, while *A*_1_ only includes cells affected by drug 1, but not drug 2 (*i.e*., 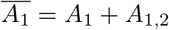). Then

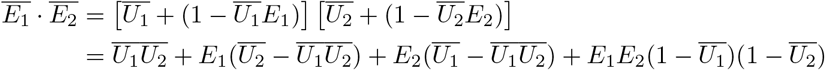

From 25, we know 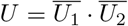, and 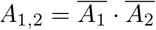, leading to

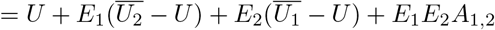

Similarly, it is simple to show 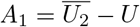, and similarly for *A*_2_

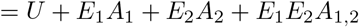

If *E*_3_ = *E*_1_ · *E*_2_, then this is equivalent to equation (26). Therefore, given *α*_12_ = *α*_21_ = 1, *γ*_12_ = *γ*_21_ = 1, *E*_0_ = 1, and *E*_3_ = *E*_1_ · *E*_2_, MuSyC predicts 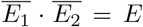. Thus, while Bliss was derived purely within the scope of “percent affected”, MuSyC shows that the Bliss model may be appropriately extended to any measure of effect for which *E*_0_ = 1 and effects are expected to be multiplicative. Nevertheless, for effects which do not satisfy these criteria, the Bliss model cannot be reliably used, while MuSyC may still be used for arbitrary effects.

#### 2.2 Dose Equivalency Principle

The DEP defines asserts an expectation that for a given effect *E*, achievable either by dose *d*_1_ of Drug 1 alone, or dose *d*_2_ of Drug 2 alone, there is a constant ratio 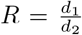 such that using Δ*d*_2_ less of Drug 2 can always be compensated for by using Δ*d*_1_ = *R*Δ*d*_2_ more of Drug 1 to achieve the same effect [11]. This definition leads to the linear isoboles characteristic of the Loewe null model.

Chou and Talalay showed that linear isoboles emerge when the two drugs are mutually exclusive [40], meaning that the double-drugged state (*A*_1,2_ in Figure 1B) is unreachable. In MuSyC, this requires setting *α*_12_ = *α*_21_ = 0, which reduces the 2D Hill equation (eq. (8)) to

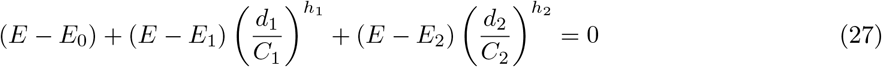

From this equation it is easy to see when *h*_1_ = *h*_2_ = 1, the conic section reduces to a line, resulting in the canonical linear isoboles of Loewe Additivity and the CI null models. Further from equation (27), we find the slope of isoble is equal to 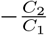 as shown by:

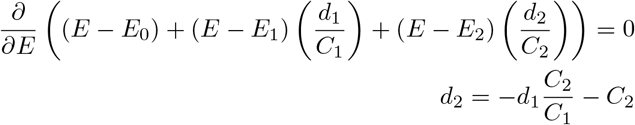

Therefore the constant *R* in the statement of the Dose Equivalence Principle is revealed by MuSyC to be equal to the ratio of the two drugs’ EC50. There is no dependence on *β* or *γ* because those parameters relate to the *A*_1,2_ state, which is blocked here. For fixed values of *E*, equation (27) results in linear isoboles only when *h*_1_ = *h*_2_ = 1. Thus, given these conditions on α and *h*, MuSyC reproduces the DEP. However, when *h* ≠ 1 nonlinear isoboles result (Figure S2), suggesting that DEP is an inappropriate expectation for such drugs (see Results Section *Re-examining the sham experiment: Sham compliance introduces Hill-dependent bias in DEPmodels* for further investigation into this issue).

### 3 Relationships between different synergy frameworks

#### 3.1 Effective dose model (Zimmer *et. al*.)

Zimmer *et. al*. [8] introduced the effective dose model as a parameterized extension of Bliss, and to our knowledge were the first to account the asymmetric potency synergy, which is also present in MuSyC. The effective dose model is constructed by fitting the dose response of each single drug to a two-parameter 1D Hill equation in which *E*_0_ and *E_max_* are fixed at 1 and 0, respectively

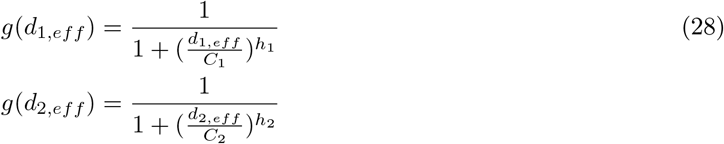

To model combination synergy, the authors propose transforming the doses *d_i_* to “effective doses” via a system of equations coupling effective doses to one another via a Michaelis-Menten term in the denominator scaled by a synergy parameter *a*.

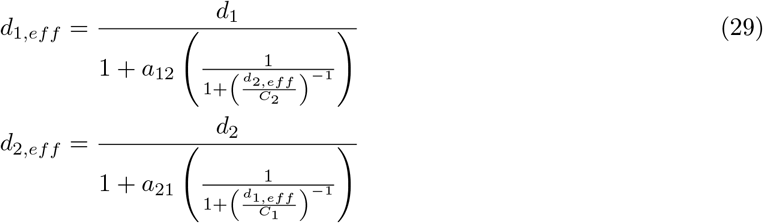

**Table S1:** Translating the effective dose model. Where possible, the original variable names from Zimmer *et. al*. have been translated to the equivalent variable names used in this manuscript, for ease of readability.

| Original variable name | MuSyC variable name |
| --- | --- |
| $n_1$ | $h_1$ |
| $D0_1$ | $C_1$ |
| $n_2$ | $h_2$ |
| $D0_2$ | $C_2$ |

The parameter *a*_12_ represents how drug 2 modifies the effective dose synergistically (*a*_12_ < 0) or antagonistically (*a*_12_ > 0) drug 1. Note that as 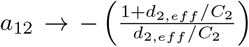, *d*_1,*eff*_ → +∞, and as *a*_12_ → +∞, *d*_1,*eff*_ → 0, which defines the bounds over which *a*_12_ is defined. The authors then fit the *a* parameters using a surface model based on MSP

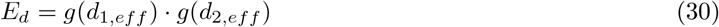

Thus the Effective Dose model reduces to the Bliss null model when *a*_12_ = *a*_21_ = 0. There are obvious similarities between Effective Dose model’s *a* parameters and MuSyC’s *α* values, as both reflect a potency transformation; however, the exact details are slightly different. For example, Zimmer assumes each drug has a Michaelis-Menten like effect on the potency of the other drugs (eq. (29)), whereas MuSyC can account for non-Michaelis-Menten effects (when *h* ≠ 1). Furthermore, by using equation 28, Zimmer explicitly assumes the measured drug effect ranges from 100% to 0%, and fit the data with this constraint. Their model is unable to accurately describe combinations where the two drugs either have unequal maximum effects, or the combination has a greater effect than the drugs can achieve alone, features which are commonly observed [41] (Figure S5). In contrast, MuSyC is able to fit dose response surfaces with arbitrary effect ranges.

#### 3.2 ZIP

In contrast to the Effective Dose Model, ZIP, accounts for changes in both the Hill slope and the potency across the dose-response surface. In ZIP, these changes are integrated into a single number (*δ*), given by

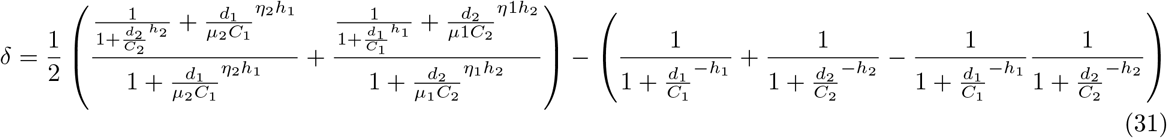

ZIP is formulated for arbitrary *E*_0_ and *E_max_*; however, it assumes *E_max_* is the same for both drugs, as well as the combination (*E*_1_ = *E*_2_ = *E*_3_). To calculate *δ*, the ZIP method fixes the concentration of one drug, then fits a Hill-equation dose response for the other drug. However, for combinations with efficacy synergy or antagonism, slices of the dose-response surface can have non-Hill, and even non-monotonic shapes. In these cases, ZIP parameter fits may not be meaningful. Because MuSyC accounts explicitly for efficacy synergy, its surfaces are able to describe such complex drug combination surfaces where ZIP cannot.

Nevertheless, ZIP parameters *μ* and *η* are closely related to MuSyC parameters *α* and *γ*. In the absence of synergistic efficacy, slices of MuSyC dose-response surfaces are sigmoidal, though in general do not perfectly follow a Hill equation, and so the ZIP model is still not identical to MuSyC. However, at saturating concentrations of one or the other drug, MuSyC does reduce to the Hill equation. In these saturating cases, ZIP’s *δ* can be related analytically to MuSyC’s *α* and *γ* by

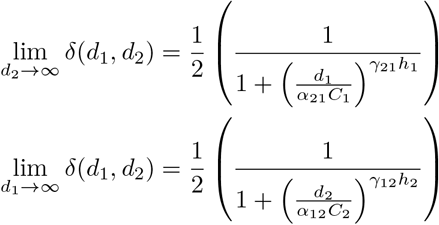

**Table S2:** Translating ZIP. Where possible, the original variable names from Yadav *et. al*. [9] have been translated to the equivalent variable names used in this manuscript, for ease of readability. Note we found an erratum flipping *x*_2_ and *m*_2_ in the step from equation 13 to equation 14 in Yadav *et. al*. which is propagated through to equation 19. Our analysis uses the intended form.

| Original variable name | Translated variable name |
| --- | --- |
| $m_1$ | $C_1$ |
| $m_2$ | $C_2$ |
| $m_{1 \rightarrow 2}$ | $\mu_1 \cdot C_2$ |
| $m_{2 \rightarrow 1}$ | $\mu_2 \cdot C_1$ |
| $\lambda_1$ | $h_1$ |
| $\lambda_2$ | $h_2$ |
| $\lambda_{1 \rightarrow 2}$ | $\eta_1 \cdot h_2$ |
| $\lambda_{2 \rightarrow 1}$ | $\eta_2 \cdot h_1$ |
| $x_1$ | $d_1$ |
| $x_2$ | $d_2$ |
| $\delta$ | $\delta$ |

#### 3.3 BRAID

BRAID [13] is an extension of the DEP to effects exceeding the weaker drug and consequently reduces to Loewe under particular conditions (Figure 2). The authors propose three BRAID models with increasing complexity, with eBRAID capable of describing the most general dose-interaction surfaces. We focus our analysis on eBRAID, which assumes that each drug alone has a Hill-like response, and constructs an Hill-like equation for the combination

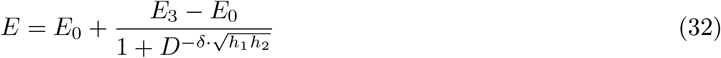

where

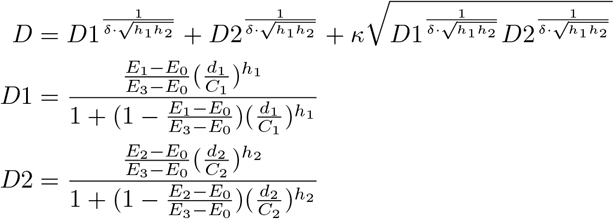

The BRAID equation (eq. (32)) uses a dose parameter, which combines the doses of both individual drugs, using a parameter *κ* and a parameter for the Hill slopes *δ* which acts as a multiplicative of the geometric mean hill slope 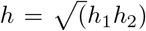. This formalism, like Loewe, is sham compliant under certain conditions, namely when 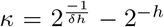. By adjusting *κ*, BRAID is able to fit complex drug combination surfaces, including non-monotonic responses, unlike ZIP. Additionally, because BRAID fits the whole combination surface using a single parameter, it can be used to make unambiguous statements about whether the combination is synergistic or antagonistic. Nevertheless, BRAID does not account for differences in synergy due to efficacy, potency, and cooperativity, whereas we find many combinations that are synergistic with respect to one, but antagonistic with respect to the other (Figure 3C). Though *κ* and *δ* are related to the potency and cooperativity respectively, their biochemical interpretation is not straightforward.

**Table S3:** Translating the BRAID model. Based on eBRAID model set of equations in the supplement. Note we corrected a typo in equation for 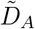 where *ID_M, B_* is suppose to be *ID_M, A_*

| Original variable name | Translated variable name |
| --- | --- |
| $D_A$ | $d_1$ |
| $D_B$ | $d_2$ |
| $\tilde{D}_{AB}$ | $D$ |
| $\tilde{D}_A$ | $D1$ |
| $\tilde{D}_B$ | $D2$ |
| $E_0$ | $E_0$ |
| $E_f$ | $E_3$ |
| $E_{f,A}$ | $E_1$ |
| $E_{f,B}$ | $E_2$ |
| $E_{AB}$ | $E_d$ |
| $n_a$ | $h_1$ |
| $n_b$ | $h_2$ |
| $ID_{M,A}$ | $C_1$ |
| $ID_{M,B}$ | $C_2$ |
| $\delta$ | $\delta$ |
| $\kappa$ | $\kappa$ |

#### 3.4 Highest Single Agent

Highest single agent (HSA) [11] is a parsimonious model that defines synergy as the net difference between the combination response and the stronger single-drug response

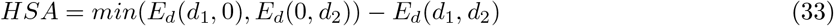

This form assumes that drug decreases *E*, though it can also be defined for drugs that increase *E*. At high concentrations of *d*_1_ and *d*_2_, equation (33) becomes proportional to our definition of efficacy synergy (*β*) as shown in equation 34 and Figure 2. Nevertheless, at intermediate doses, HSA will conflate synergy of potency, efficacy, and cooperativity (Figure 3) highlighting the importance of considering the whole dose-response surface when calculating synergy.

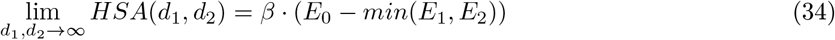

#### 3.5 2D Hill PDE

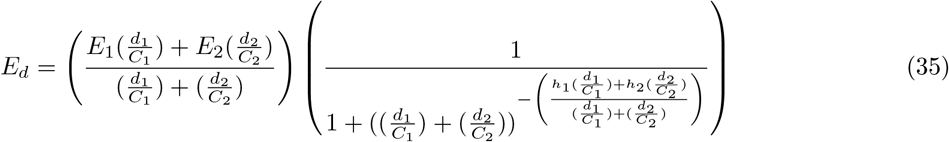

The most recent framework is one by Schindler [10] which interpolates a null dose-response surface from the single dose-response curves alone without any fit parameters. This was done by using PDE Hill equations and then imposing boundary conditions as well as sham compliance. It is therefore an extension of Loewe to effects greater than the least efficacious drug (Figure 2, Table 2). The boundary conditions enforce the null model’s maximal effect of the combination (*E*_3_) is equal to the mean of *E*_1_ and *E*_2_. This results in the non-intuitive scenarios such as if the maximal effect of one compound is 0.25 and the other is 0.75, then the predicted maximal effect of the combination is 0.5 which is much less than achievable with a single drug.

**Table S4:**
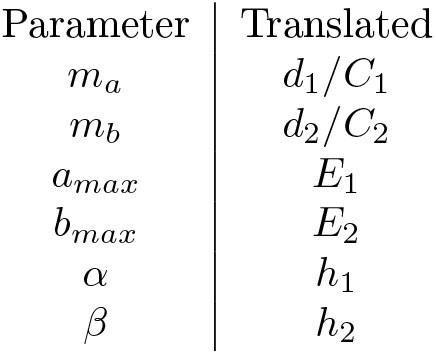
Translating the Schindler *et al*. model. Based on equations 7 and 8 in the main text.

| Parameter | Translated |
| --- | --- |
| $m_a$ | $d_1/C_1$ |
| $m_b$ | $d_2/C_2$ |
| $a_{max}$ | $E_1$ |
| $b_{max}$ | $E_2$ |
| $\alpha$ | $h_1$ |
| $\beta$ | $h_2$ |

### 4 Sham Compliance of Synergy Frameworks

To verify a new synergy model’s consistency, it is traditionally tested with the “sham” combination thought experiment. Briefly, the thought experiment proposes a single drug is divided into two vials labeled drugs “A” and “B”, before the vials are given to an unsuspecting researcher —who does not know they are the same drug —to perform combination synergy measurements [3]. Any synergy metric finding either synergy or antagonism for this “sham combination” fails the thought experiment, because a drug combined with itself should be additive. HSA, as well as Bliss and other MSP frameworks (Figure 2) famously fail the sham experiment [11]. In contrast, Loewe additivity and other DEP frameworks are sham complaint.

In reviewing the literature, we identified a two errors beyond the Hill slope bias (Figure 5) pertaining to the sham experiment that merit addressing.

1. Chou, one of the creators of the Combination Index (CI), has strongly argued that satisfaction of the sham experiment is critical. However, we identified an error in the derivation of the CI from its underlying model, so that while the CI equation is sham-compliant, the biochemical model proposed by Chou and Talalay is not.
2. ZIP was proposed as a framework that unifies Loewe Additivity (DEP) and Bliss Independence (MSP), much like MuSyC. In support of this, they prove ZIP is sham-compliant; however, we identified an error in their proof. Further, we generate *in silico* sham response data and show that ZIP fails the sham experiment. For this reason, we place ZIP as an MSP method in Figure 2C, and not DEP. However, we have contended that satisfying the sham experiment should not be a sought-after standard in drug combinations, so we do not consider it a shortcoming of ZIP that it does not.

#### 4.1 Sham Compliance of Combination Index

Chou and Talalay report that the combination index for mutually exclusive drugs satisfies the sham experiment, whereas for MuSyC, we find this is only true when *h* = 1 (eq (10)). Because MuSyC’s underlying model (Figure 2B) is identical to what Chou and Talalay describe [40] when *α*_12_ = *α*_21_ = 0, we sought to discover the source of this discrepancy.

Chou and Talalay’s finding comes from “Generalized Equations for the Analysis of Inhibitions of Michaelis-Menten and Higher-Order Kinetic Systems with Two or More Mutually Exclusive and Nonexclusive Inhibitors” [40]. We found an error in the section *Inhibition of the Higher-Order Kinetic Systems by Mutually Exclusive Inhibitors*. Specifically, the authors correctly solve the n-drug case with *h* =1

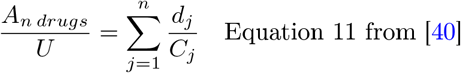

as well as the the 1-drug case with arbitrary *h*

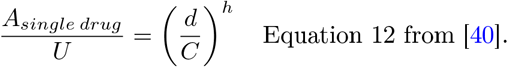

However, they incorrectly extrapolate these two equations to an *n*-drug, arbitrary *h* case (though they assume all drugs have the same value of *h, e.g*. all drugs have *h* binding sites and follow Hill-type kinetics) by erroneously stating

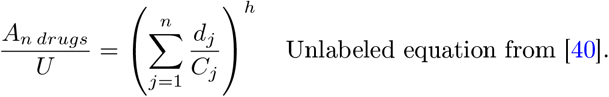

In general it is not clear how this would be calculated if each drug has a different value *h*, as there exists only a single *h* in their formula. However, in the case of *n* mutually exclusive drugs following Hill kinetics (a model graphically represented for *n* = 2 in Figure 2B), we can say at equilibrium

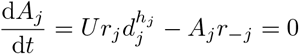

From this we can solve for 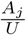 for each drug, arriving at the same equation 12 from Chou [40]. But applying the constraint that *U + A* = 1, where 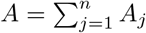, we instead find the correct form for the multi-drug case with arbitrary *h* is

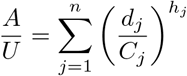

When *h_j_* = *h* = 1 the discrepancy between these equations vanishes because it does not matter that they place the exponent outside the sum. However Chou and Talalay use the incorrect version to define their Combination Index [25], regardless of *h*.

#### 4.2 Sham Compliance of ZIP

ZIP was reported to satisfy the sham experiment [9] based on an argument that if both drugs are the same, then *m*_1_ = *m*_2_ = *m*_1←2_ = *m*_2←1_. However, this statement is false for sham experiments, because once some of the drug is added, *m*_1←2_ is shifted. To demonstrate this, consider their model (equation 13 from [9], without asserting *m*_1←2_ = *m*_1_)

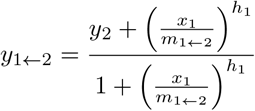

*m*_1←2_ is the amount *x*_1_ of drug 1 that is needed to achieve an effect halfway between *y*_2_ and 1 (the asymptotic value at infinite drug), where *y*_2_ is the effect achieved by adding an amount *x*_2_ of drug 2. Specifically, when 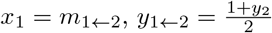.

Sham experiments satisfy Equation (11), which combined with the above gives

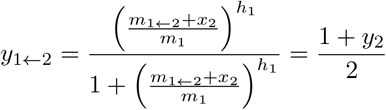

Further we know

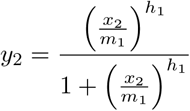

This system can be solved to find

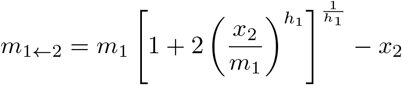

demonstrating that *m*_1←2_ ≠ *m*_1_ for the sham experiment in the ZIP model, contradicting their proof that ZIP is sham-compliant.

To verify ZIP identifies synergy or antagonism for sham combinations, we generated a synthetic sham dose response surface. Sham experiments can be generated exactly for any drug with a pre-defined dose-response by asserting the condition in Equation (11). We constructed a synthetic dataset describing a sham dose response surface for a drug with *h* = 2, sampled at 2.5 orders of magnitude above and below the EC50. One drug was sampled at 7 concentrations, the other at 12, defining a 7 x 12 sham dose response matrix. We used the synergyfinder [16] R package to calculate synergy by both Loewe and ZIP (Figure S6), and found Loewe reported close to 0 synergy, as expected for a sham combination, but confirms ZIP detects synergy and antagonism at several concentrations.

### 5 Percent Affect vs Percent Effect

It is generally known that Bliss, and more generally the MSP methods, are only applicable to percentage data. Indeed, percent transformations are commonly applied to data in order to apply Bliss. In Bliss’ original study, drug effect was quantified as the percentage of eggs killed (the probability that each egg would die at a given toxin dose), but in all cases the measurement was a discrete event (death of an insect egg). Therefore, the metric of drug *effect* was the a percentage of affected eggs. However, to apply Bliss to more general measures of drug effect for which discrete counts cannot be obtained, it has been ubiquitous practice to normalize the drug effect as a percent relative to control (*e.g*., percent viability in cancer research). Nevertheless, such normalization does not, in general, transform measures of efficacy into measures of percent affect.

Suppose, as an example, analyzing a drug treatment which actually caused the cells to grow slightly faster than control. By normalizing the drug effect to control, the percent viability is greater than 100% which cannot mean that >100% of the cells were *affected*.

Alternatively, consider the case when a cytostatic drug causes all treated cells to halt both proliferation and death. If the control population doubled twice over 72 hours, the percent viability would be 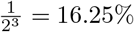. If the measure of percent viability was taken instead at 96 hours, the percent viability would be 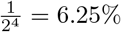. At both time points the percent of affected cells was the same (100%); however, the percent of drug *effect* changes due to normalization.

### 6 Proof of boundary behavior of the 2D Hill equation

When *d*_1_ → ∞, then the 4 state reduces to 2 states transition model between A1 and A12.

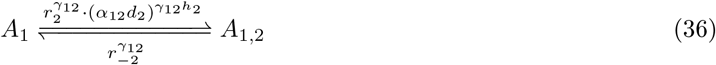

At equilibrium

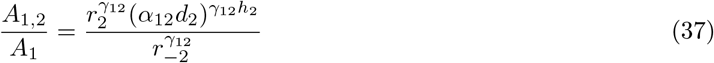

When *A*_1_ = *A*_1,2_ this is the EC50 for drug 2 given saturating concentrations of drug 1 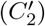. This dose can be found by solving the above.

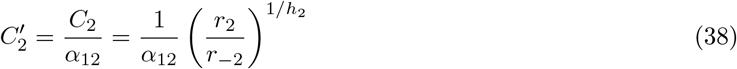

The boundary then reduces to a Hill equation of the form

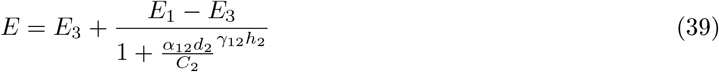

following the derivation in box 1.

